# HP-Bodies – Ancestral Condensates that Regulate RNA Turnover and Protein Translation in Bacteria

**DOI:** 10.1101/2025.02.06.636932

**Authors:** Jian Guan, Rebecca Lee Hurto, Akash Rai, Christopher A. Azaldegui, Luis A. Ortiz-Rodríguez, Julie S. Biteen, Lydia Freddolino, Ursula Jakob

**Affiliations:** Department of Molecular, Cellular and Developmental Biology, University of Michigan, Ann Arbor, MI, USA; Department of Biological Chemistry, University of Michigan, Ann Arbor, MI, USA; Program in Chemical Biology, University of Michigan, Ann Arbor, MI, USA; Department of Chemistry, University of Michigan, Ann Arbor, MI, USA; Department of Computational Medicine and Bioinformatics, Ann Arbor, MI, USA

**Keywords:** Biological Condensates, Stress, Processing Bodies, Bacteria, RNA stability, Hfq, Polyphosphate

## Abstract

Uncovering what drives select biomolecules to form phase-separated condensates *in vivo* and identifying their physiological significance are topics of fundamental importance. Here we show that nitrogen-starved *Escherichia coli* produce long-chain polyphosphates, which scaffold the RNA chaperone Hfq into phase-separating high molecular weight complexes together with components of the RNA translation and processing machinery. The presence of polyphosphate within these condensates, which we termed HP-bodies, controls Hfq function by selectively stabilizing polyadenylated RNAs involved in transcription and protein translation, and promoting interactions with translation- and RNA-metabolism-associated proteins involved in *de novo* protein synthesis. Lack of polyphosphate prevents HP-body formation, which increases cell death and significantly hinders recovery from N-starvation. In functional analogy, we demonstrate that polyP contributes specifically to the formation of Processing (P)-bodies in human cell lines, revealing that a single, highly conserved and ancestral polyanion serves as the universal scaffold for functional phase-separated condensate formation across the tree of life.

## INTRODUCTION

Membrane-less organelles, also referred to as biomolecular condensates, arise from a process known as liquid-liquid phase separation (LLPS), where biomolecules in solution separate into a condensed liquid phase surrounded by a dilute phase ^1,2^. Condensates serve as a critical addition to the endomembrane system in the subcellular organization of diverse physiological processes as they promote the specific segregation of biochemical pathways and local enrichment of cellular components^3^. In the vast majority of prokaryotes, the endomembrane system is entirely absent ^4^, potentially increasing the importance of LLPS for compartmentalizing biological processes. However, in stark contrast to eukaryotic organisms where biomolecular condensates such as P-bodies and stress granules have been extensively studied ^5–7^, little is known about their roles in bacteria ^4^. At present, the best characterized bacterial condensates are <u>b</u>acterial <u>R</u>NP (<u>BR</u>)-bodies in *Caulobacter crescentus*^8–10^. Scaffolded by the intrinsically disordered region (IDR) of RNase E, BR-bodies recruit long poorly translated mRNAs for degradation, thereby reshaping the bacterial transcriptome to cope with stress ^9^. Recent studies in *E. coli* revealed that the conserved RNA chaperone Hfq forms condensate-like foci in response to either N-starvation or osmotic stress ^11–13^. Intriguingly, Hfq is a small Sm-like protein that shares structural features with eukaryotic Sm/LSm proteins, the scaffolding components of P-bodies ^14^. Like other Sm-like proteins, the N-terminus of Hfq assembles into a toroidal hexamer core, whereas the C-terminus of *E. coli* Hfq is intrinsically disordered ^15^. The physicochemical heterogeneity across the hexameric surface results in multiple RNA/DNA binding interfaces with distinct specificities ^16^. Best known for its function as a RNA chaperone, Hfq facilitates sRNA-mRNA duplex formation, thus regulating mRNA stability and translation ^17,18^. In addition, at least 30% of cellular Hfq is nucleoid-associated and involved in the silencing of chromosomal regions that are enriched for prophages and other mobile genetic elements ^19–21^. In response to some stress conditions (most notably nitrogen (N)-starvation and hyper-osmotic shock), Hfq assembles into a single, polarly-localized condensate together with select sRNAs, RNase E and other RNA degradosome components ^11–13,22^. Yet very little is known regarding why and how Hfq forms a single condensate during these *in vivo* stress conditions, what other proteins, RNA, and/or DNA components are involved, and how Hfq condensate generation contributes to stress survival. Given the known involvement of polyanions such as RNAs in condensate formation ^23–25^, we investigated whether polyphosphate (polyP) — the most ancient, ubiquitous, and highly negatively charged polymer known — might constitute a critical component of these stress-induced Hfq condensates ^26^. We based this reasoning on previous reports, which demonstrated that i) the stress conditions where Hfq condensates are observed also trigger polyP accumulation ^27^, ii) polyP forms distinct granules in N-starved *Pseudomonas aeruginosa* ^28,29^, and, most critically, iii) that polyP phase separates with Hfq *in vitro* ^21^. Indeed, we now demonstrate that long-chain polyP is crucially important to organize Hfq condensate formation *in vivo*. We find that the Hfq-polyP condensate specifically accumulates and selectively stabilizes polyadenylated transcripts that are critical for bacterial survival under and recovery from nitrogen-starvation. Extending to the extreme opposite end of the tree of life, we realized that polyP accumulation is also required for the formation of mammalian P-bodies but not stress granules, indicating an evolutionarily conserved pathway for membrane-less organelle formation. Taken together, our findings demonstrate that polyP plays a conserved role in shaping functional biomolecular condensates throughout evolution, marking it as perhaps the most ancient and conserved scaffold for condensate formation across biology.

## RESULTS

### PolyP drives Hfq foci formation in *E. coli*

To investigate the potential influence of polyP on Hfq foci formation *in vivo*, we replaced the chromosomal copy of *hfq* with a fully functional *hfq-mCherry* gene fusion ^30^ in either *E. coli* MG1655 wild type (WT) or a mutant strain that lacks polyphosphate kinase (PPK), the only known polyP-synthesizing enzyme in *E. coli* ^31^. When grown in defined media supplemented with limiting amounts of nitrogen (N), both strains enter N-starvation at ∼5 h (N0) (Fig. S1A). At this point, WT bacteria start to accumulate polyP in the form of very long (>300 P_i_ units) chains (i.e., polyP-300) (Fig. 1A) and begin to display a single Hfq focus at one of the cell poles (Fig. 1B). Within 24 hours of nitrogen-starvation (henceforth, N24), over 80% of all WT cells contain one polarly localized Hfq focus (Fig. 1C). In stark contrast, the *ppk* deletion mutant strain, which does not produce any detectable polyP (Fig. 1A), shows a substantial delay in the appearance of Hfq foci with fewer than 40% of the mutant bacteria containing visible Hfq foci at N24 (Fig. 1B, C). This result was surprising given that the N24 *ppk* deletion mutant accumulates almost twice the amount of total Hfq compared to N24 WT *E. coli* (Fig. S1B). We also observed that most focus-containing *ppk* bacteria harbor multiple and smaller Hfq foci (Fig. 1B, S1C), which are less frequently enriched at the poles (Fig. S1D). We obtained similar results when we tagged Hfq with a 3×FLAG tag and visualized Hfq localization in fixed cells, thereby excluding the possibility of mCherry-induced artifacts (Figure S1E). Importantly, we also reproduced these results in osmotically stressed WT *E. coli* cells: upon treatment with NaCl for 4 h, WT bacteria show a substantial increase in the levels of long and medium-chain length polyP (Fig. 1D) as well as display a single, clearly visible and polarly localized Hfq focus (Fig. 1E). In contrast to the ∼80% of focus-containing WT bacteria, fewer than 5% of the *ppk* bacteria display any detectable focal accumulation of Hfq under hyperosmotic stress conditions (Fig. 1E, F). Based on these results, we now wondered whether the accumulation of polyP might serve as the actual driving force for Hfq focus formation. To test this idea, we artificially increased the cellular polyP levels in the *ppk* deletion strain by expressing the mutant variant *ppk10*, previously found to substantially increase polyP synthesis in exponentially growing *E. coli* cells above WT levels^32^ (Fig. 1G). Indeed, we observed that this increase in polyP levels was sufficient to cause the formation of Hfq foci even in the absence of additional stress (Fig. 1H), demonstrating that stress-induced increases in cellular polyP concentrations play a critical regulatory role in coalescing Hfq into distinct foci.

**Figure 1.**
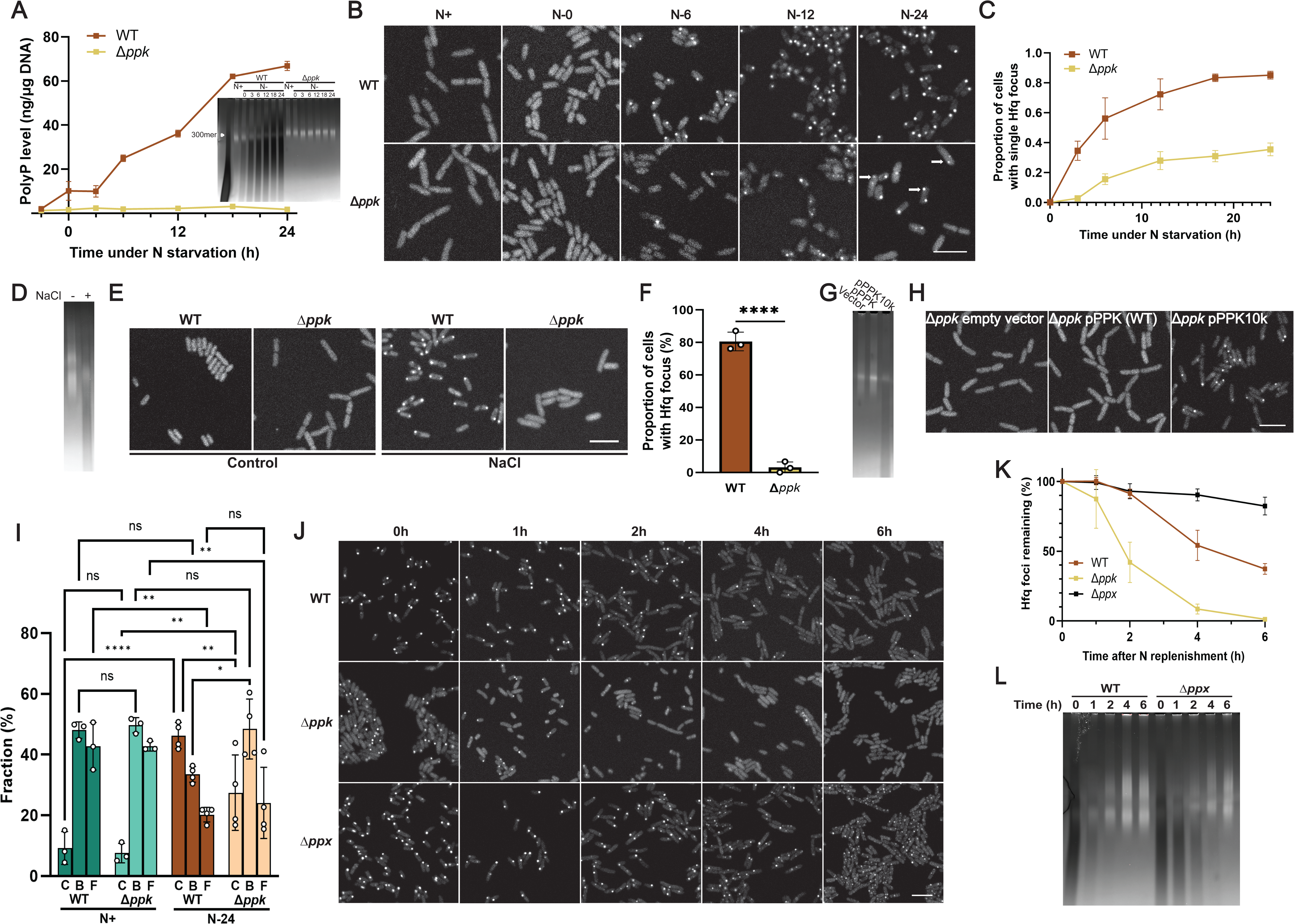
PolyP drives foci formation of Hfq in *E. coli*. **(A)** Normalized polyP levels in WT and Δ*ppk E. coli* 2 hours before and select timepoints upon entry into N-starvation (N0). Error bars indicate SD (n = 3). **Inset:** PolyP extracted under the same conditions and visualized on TBE gel by DAPI staining and photobleaching, which will appear bright for nucleic acid and dark for polyP. Purified polyP 300mer was used as reference (lane 1). **(B)** Subcellular localization of Hfq-mCherry in WT or Δ*ppk E. coli* MG1655 *hfq::hfq-mCherry* during exponential growth (N+) or at select times after onset of N-starvation (N-). Arrows indicate cells with multiple smaller Hfq foci. **(C)** Quantification of cells containing visible Hfq-mCherry foci after entering N-starvation. A minimum of 200 cells were quantified for each of three replicates under each time point. Error bars show SD. **(D)** PolyP extracted from *E. coli* MG1655 *hfq::hfq-mCherry* after 4 hours of mock treatment (-) or upon osmotic stress treatment with 1.17 M NaCl (+) was visualized on TBE gel by DAPI staining and subsequent photobleaching. **(E)** Subcellular localization of Hfq-mCherry in MG1655 *hfq::hfq-mCherry* WT or Δ*ppk* after 4 hours of osmotic stress treatment. **(F)** Quantification of cells containing visible Hfq-mCherry foci after osmotic stress. At least 100 cells were quantified in each of three replicates. Comparisons were made by unpaired *t*-tests. ****p<0.0001. **(G)** PolyP extracted from (N+) Δ*ppk E. coli hfq::hfq-mCherry* carrying an empty plasmid or a plasmid overexpressing WT *ppk* or *ppk10k* was visualized on TBE gel by DAPI staining and subsequent photobleaching, which darkens the polyP signal. **(H)** Subcellular localization of Hfq-mCherry in (N+) Δ*ppk E. coli hfq::hfq-mCherry* carrying an empty plasmid or a plasmid overexpressing WT *ppk* or *ppk10k*. Scale bars represent 5 µm. **I)** Fraction of PAmCherry Hfq present in condensates (C), found in the nucleoid-bound (B) form or free (F) in the cytosol of N+ and N24 WT or *ppk* deletion strains as assessed by single-molecule localization. > 1500 trajectories were analyzed per replicate (*N* ≥ 3) of each condition. Comparisons were made by 2-way ANOVA (n ≥ 3), *p<0.05, **p<0.01, ****p<0.0001. Error bars indicate SD. **(J)** Subcellular localization of Hfq-mCherry in N24 WT, Δ*ppk,* or Δ*ppx E. coli* MG1655 *hfq::hfq-mCherry* at indicated time points after supplementing the media with fresh nitrogen. **(K)** Percentage of Hfq-mCherry foci remaining calculated from (B). The number of condensates at N24 (i.e., t=0h) is set to 100 % and used as a reference point. A minimum of 200 cells were quantified for each of three replicates under each time point, error bars indicate SD. **(I)** PolyP levels at the indicated timepoints after supplementing the growth media of N24 MG1655 *hfq::hfq-mCherry* WT or Δ*ppx* with fresh N-source. PolyP was visualized on a TBE gel using DAPI staining and subsequent photobleaching, which darkens the polyP signal.

### PolyP controls the cellular reorganization of Hfq during and after N-starvation

To investigate the effects of N-starvation on the localization and dynamics of Hfq in more detail, we used single-molecule super-resolution microscopy and compared the mobility of Hfq-mCherry molecules in WT and *ppk* deletion mutants grown under N-replete (N+) or N-starvation conditions (N24). Based on Hfq’s diffusion rate and localization relative to the observed foci (see Methods for details), we were able to assign each Hfq molecule to either inclusion in a slow-diffusing condensate (C), slow-moving nucleoid-bound state (B) or fast-diffusing free state (F) (Fig. 1I). Consistent with the bulk observations noted above, we found that about 46% of all Hfq molecules in N24 WT *E. coli* coalesce into one distinct slow-diffusing C state (i.e., the Hfq focus) while only ∼27% of Hfq molecules are associated in the C state in the N24 *ppk* deletion mutant (Fig. 1I). Furthermore, we noted that in WT bacteria, the Hfq molecules that form the slow-diffusing C state are recruited from both the pool of highly mobile “free” (F) Hfq molecules (∼23%) and, to a lesser extent (∼14%), from nucleoid-bound (B) Hfq molecules (Fig. 1I). In the absence of polyP, however, Hfq appears to be primarily recruited from the free pool, thus leaving close to 50% of non-focus associated Hfq molecules bound to the nucleoid. These results agree well with previous *in vitro* findings, which showed that, at sufficiently high concentrations, polyP competes with DNA for Hfq binding ^21^. We thus infer that in addition to directly interacting with Hfq to mediate focus/condensate formation, polyP plays the additional role of selectively releasing nucleoid-bound Hfq (through competition with DNA binding) to free additional Hfq hexamers for focus formation. To test whether polyP also affects the reverse reaction (i.e., focus dispersal), we then treated N24 WT and *ppk* deletion mutants with fresh nitrogen (N), which is known to cause the dissolution of Hfq foci within a few hours ^12^. Indeed, we observed that the Hfq foci disappear from WT *E. coli* at a rate that coincides with the decline in cellular polyP levels (compare Fig. 1J, K with Fig. 1L). In contrast, Hfq foci formed in the *ppk* deletion mutant disappear significantly faster than in WT *E. coli*. Furthermore, foci that formed in an *E. coli* mutant that lacks the polyphosphatase (PPX)—and hence harbors increased polyP levels during stress and shows a slower polyP decline upon recovery (Fig. 1L)—reveal a significant delay in their disappearance (Fig. 1J, K). These results clearly demonstrate that changes in the cellular levels of polyP affect the formation as well as the dispersal of Hfq foci by partitioning Hfq molecules across various cellular compartments (cytoplasm, condensate, nucleoid).

### The scaffolding function of polyP is relevant for HP-body formation *in vivo*

To directly test whether polyP constitutes a component of the Hfq foci in N24 WT *E. coli*, we fixed Hfq-mCherry expressing WT and *ppk* mutant bacteria grown for 24h under N-starvation conditions, and co-stained the cells using an antibody against mCherry to detect Hfq and a PPX-derived polyP probe (PPXBD) to stain for polyP ^33^ (Fig. 2A). We detected a clear colocalization between the two components in N24 WT bacteria but not in the polyP-depleted *ppk* deletion mutant (Fig. 2B), further supporting the conclusion that the association of polyP with Hfq promotes foci formation. Based on these results, we reasoned that the effects of long-chain polyP on *in vivo* Hfq condensate formation might be related to the previously observed ability of long chain polyP to scaffold Hfq hexamers into higher molecular weight (HMW) assemblies *in vitro*^21^. These *in vitro* formed HMW assemblies of Hfq revealed a clear polyP chain-length dependence with the longest tested polyPs (i.e. polyP-300) forming supramolecular complexes that migrate very slowly on native PAGE. To test this idea, we prepared lysates from N0 and Hfq-mCherry expressing N24 WT and Δ*ppk* bacteria and analyzed the migration of Hfq-mCherry via native PAGE followed by western blots targeting Hfq-mCherry. As expected, we found that Hfq-mCherry migrates as low molecular weight species in lysates prepared from either N0 WT or N0 *ppk* deletion bacteria. In contrast, however, the majority of Hfq-mCherry in lysates from N24 WT *E. coli* forms HMW assemblies, similar to the ones we previously observed *in vitro*^21^ (Fig. 2C). Importantly, we did not observe a significant signal at the same HMW position when we analyzed the lysate of the *ppk* deletion mutant (Fig. 2C), suggesting that the smaller polyP-deficient Hfq foci found in the *ppk* deletion strain either fail to assemble into HMW complexes or are less stable and dissociate over the time course of the experiment. When we incubated the N24 WT lysate prior to the native PAGE with the polyP-degrading yeast polyphosphatase (yPPX)^34^, the majority of HMW complexes dissociated into low molecular weight Hfq species, whereas neither incubation with DNase I nor RNase A had a significant effect on the migration behavior of the HMW assemblies of Hfq (Fig. 2D). These experiments demonstrated the unique role for polyP in scaffolding HMW complex formation *in vivo*. To determine whether the presence of polyP in these HMW complexes of Hfq affects the material properties of Hfq foci *in vivo*, we next compared the rate and extent of fluorescence recovery after photobleaching (FRAP) in similarly sized foci formed in either N24-grown WT or *ppk* deletion cells (Fig. 2E). We noted that while the kinetics of the Hfq-mCherry fluorescence recovery were similar in both strains, proceeding on the order of tens of seconds, the relative levels of recovered Hfq-mCherry fluorescence were lower in the absence of polyP (Fig. 2F). These results suggested that polyP might improve the solubility of Hfq within the foci, and/or alter the material properties of the foci themselves. Indeed, analysis of the levels of soluble *versus* insoluble Hfq in WT and *ppk* mutant cells demonstrated that about 15% of Hfq molecules present in N24 starved *ppk* deletion mutants are insoluble, compared to the less than 5% of pelletable Hfq molecules in WT *E. coli* (Fig. 2G). Taken together, these results demonstrate that polyP plays a critical role in promoting Hfq focus formation and maintaining Hfq in a soluble and presumably functional state. To acknowledge this crucial role of polyP in the formation of functional Hfq-foci in WT *E. coli*, we decided to rename these condensates as “Hfq-PolyP bodies”, or HP-bodies.

**Figure 2.**
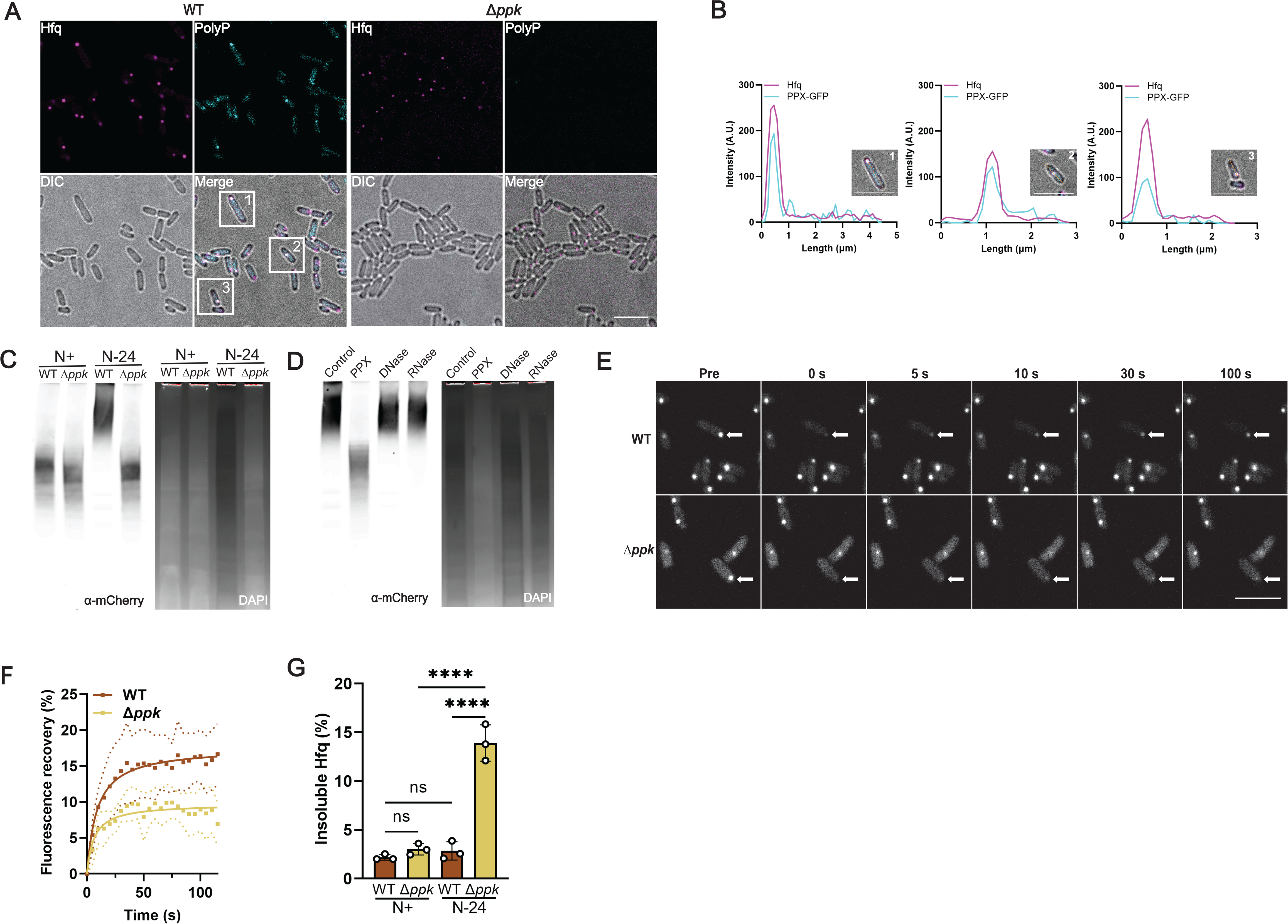
PolyP scaffolds Hfq into HMW assemblies with distinct dynamic properties. **(A)** Co-localization of polyP and Hfq-mCherry in fixed N24 WT and Δ*ppk* MG1655 *hfq::hfq-mCherry*. PolyP is visualized by immunofluorescence using a PPXBD-GFP probe while Hfq is visualized using an anti-mCherry antibody. **(B)** Plot profiles of Hfq-mCherry and PPXBD-GFP in representative cells highlighted in panel E, showing colocalization of Hfq and polyP. **(C)** Native western blot of cell lysates from MG1655 *hfq::hfq-mCherry* WT and Δ*ppk* at N+ and N24 using antibodies against mCherry. UV-bleached DAPI stain of native gel is shown in parallel. PolyP is shown as dark areas due to the behavior of the bleaching. **(D)** Native western blot of N24 in MG1655 *hfq::hfq-mCherry* WT relative to Δ*ppk* using antibodies against mCherry. Lysates were treated with the indicated enzymes for 2 hours prior to electrophoresis. UV-bleached DAPI stain of native gel is shown in parallel. **(E)** *In vivo* FRAP of similarly-sized Hfq-mCherry foci (arrow) at N24 in MG1655 *hfq::hfq-mCherry* WT and Δ*ppk* cells. **(F)** Analysis of the *in vivo* FRAP measurements shown in (G). At least 8 cells were analyzed per sample. FRAP traces were fitted against a one-phase association curve: WT _t1/2_ = 8.8 s (95% CI 6.0 s to 12.5 s), Δ*ppk* _t1/2_ = 6.2 s (95% CI 3.9 s to 9.2 s), with solid lines representing fitted FRAP curves, square dots average fluorescence intensities at indicated time points and round dotted lines indicating the SD. **(G)** Proportion of insoluble Hfq-mCherry at N0 and N24 in WT and *ppk* deletion strain as determined by quantitative western blot using antibodies against mCherry (n=3, error bars indicate SD, ****p<0.0001).

### HP-body formation increases bacterial resistance against N-starvation stress

One of the most crucial and largely unsolved questions in the field of biological condensates concerns their physiological function(s). Previous studies revealed that the deletion of *hfq*, while not affecting *E. coli* growth under exponential conditions (Fig. S1A), significantly reduces bacterial survival during N-starvation (which we recapitulated; Fig. 3A) ^12^ and delays the restart of bacterial growth upon N-resupplementation (Fig. 3B). Yet, it had been entirely unclear whether these phenotypes are connected to the overall lack of Hfq as RNA chaperone and/or nucleoid associated suppressor of gene expression, or the specific absence of free (cytoplasmic), nucleoid-bound, or HP-body associated Hfq species. Based on our discovery that the N24 *ppk* deletion strain accumulates almost twice as much total Hfq compared to the N24 WT strain (Fig. S1B), the vast majority of which are in either the free or nucleoid-bound form (Fig. 1I), we now reasoned that the phenotypical analysis of the *ppk* mutant should shed light into the physiological relevance of HP-body formation during N-starvation. We found that while the deletion of *ppk* causes a somewhat weaker phenotype in regard to N-starvation induced cell death compared to the *hfq* deletion mutant (Fig. 3A), both strains show an equally pronounced delay in the recovery upon N-resupplementation (Fig. 3B). Moreover, and even more importantly, we discovered that the combined deletion of both *ppk* and *hfq* genes precisely phenocopies the *hfq* deletion strain, indicative of an epistatic relationship between polyP and Hfq (Fig. 3A, B). These results clearly indicate that polyP and Hfq are part of the same pathway that protects *E. coli* against N-starvation stress and provide evidence for a model in which polyP acts through or upstream of Hfq. Even more importantly, however, the results provide strong evidence for the crucial role that the formation of HP-body plays for bacterial survival during and recovery from N-starvation. In addition, we discovered that whereas plasmid-borne *ppk* overexpression specifically rescued the phenotype of the *ppk* mutant only, overexpression of *hfq* rescued the phenotypes of both *hfq* and *ppk* deletion mutants (Fig. 3C, D). These results strongly suggest that the observed stress-induced accumulation of polyP lowers the critical Hfq concentration that is necessary for the formation of functional *in vivo* condensates of Hfq. Our findings also imply that the *ppk* deletion strain would be likely be even more phenotypically similar to the *hfq* deletion mutant (i.e., showing stronger defects upon N-starvation) if it were not for the apparent compensatory upregulation of Hfq (Fig. S1B).

**Figure 3.**
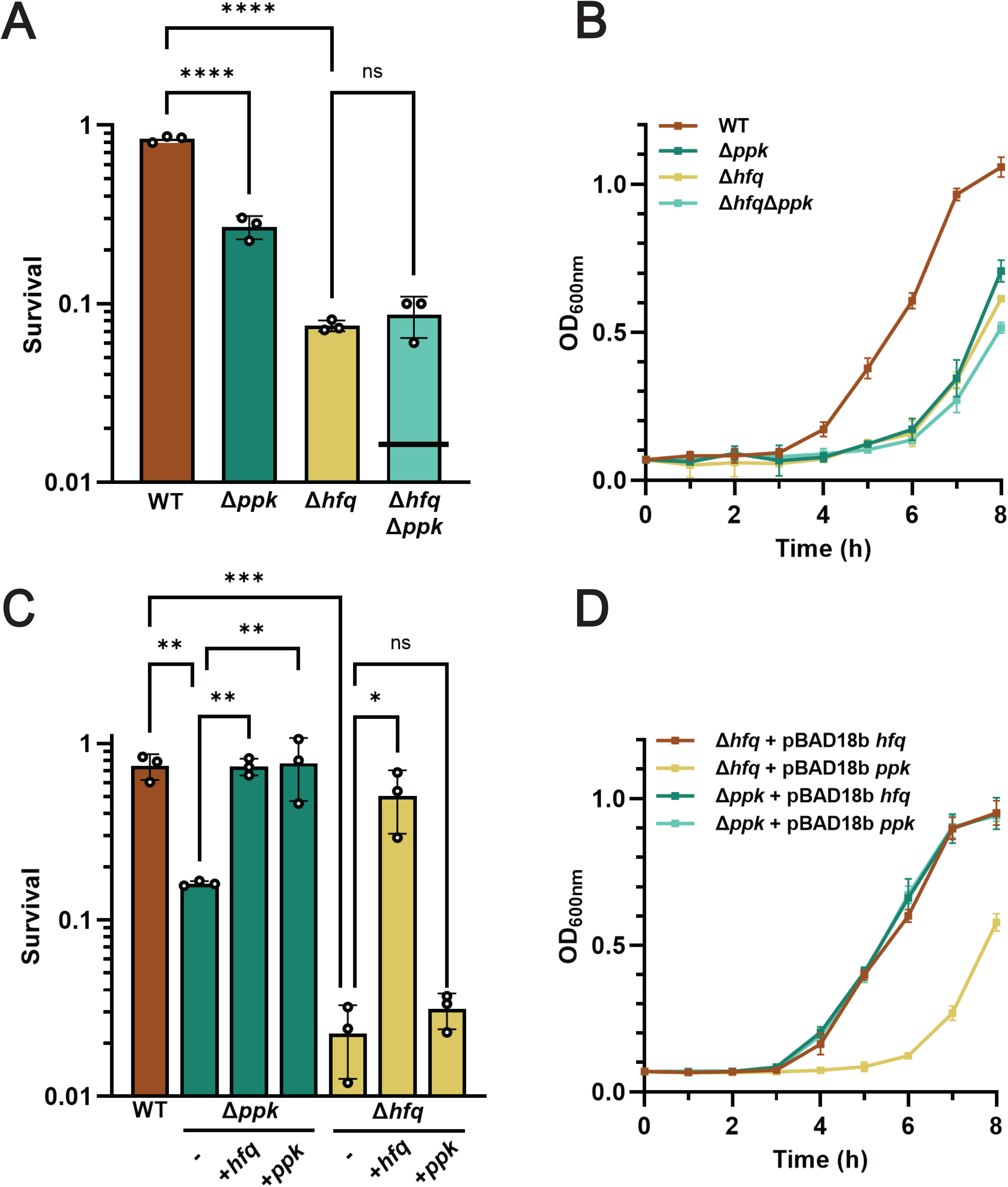
Formation of HP-bodies promotes bacterial survival during N-starvation. **(A)** Survival of WT, Δ*ppk*, Δ*hfq* and Δ*hfq*Δ*ppk E. coli* MG1655 at N48. Similar results were obtained in the respective MG1655 *hfq::hfq-mCherry* strains. The black bar in the double mutant indicates the expected phenotype if *ppk* and *hfq* deletion behaved log-additively. **(B)** Recovery growth of N24 WT, Δ*ppk*, Δ*hfq* and Δ*hfq*Δ*ppk E. coli* after dilution into fresh Gutnick medium supplemented with 3 mM NH_4_Cl. **(C)** Survival of WT and Δ*ppk* and Δ*hfq E. coli* MG1655 carrying empty vector or *ppk* or *hfq* expression vectors at N48 **(D)** Recovery growth of N24 Δ*ppk* and Δ*hfq E. coli* carrying *ppk* or *hfq* expression vectors after dilution into fresh Gutnick medium supplemented with 3 mM NH_4_Cl. Error bars indicate SD (n=3). One-way ANOVA with was used in panels A and C; *p<0.05, **p<0.01, ***p<0.001, ****p<0.0001.

### The protein composition of HP-bodies functionally resembles mammalian stress bodies

Previous studies in N-starved or osmotically stressed *E. coli* cells revealed that Hfq colocalizes with members of the RNA degradasome, including the RNA degrading enzyme RNaseE, the RNA helicase RhlB, the polyribonucleotide nucleotidyltransferase Pnp (PNPase), and enolase (Eno)^22^ . To test whether absence of polyP negatively affects these associations with Hfq, we individually tagged the members of the RNA-degradosome with mTurquoise2 at their endogenous loci in Hfq-mCherry expressing WT or *ppk* deletion strains. Live cell imaging revealed that all four members of the degradosome strongly co-localize with HP-bodies in N24 WT cells but not with Hfq foci formed in the *ppk* deletion strain (Fig. 4A). We confirmed these results by analyzing the lysates of those cells on native PAGE, which demonstrated that most of the mTurquoise2 RNase E signal associates with the HMW species of Hfq in N24 WT lysates but migrates as low molecular weight species in the *ppk* deletion strain or in PPX-treated WT lysates (Fig. 4B). To further explore the protein composition of the HMW Hfq-polyP complexes (i.e., HP-bodies), we next excised the respective gel regions and conducted MS/MS analysis (Fig. 4C). We obtained highly consistent results and identified almost 80 proteins, which are significantly enriched in the gel region corresponding to the HMW Hfq-polyP complexes in N24 WT cells but not in the corresponding gel region of the *ppk* lysate (Table S1, Supplementary Dataset 1). As expected, we observed a strong polyP-dependent enrichment for Hfq, providing further confirmation that these HMW complexes are indeed Hfq-based. Importantly, many of these proteins were depleted when we treated the WT lysates with yPPX prior to the native PAGE (Table S1), indicating that it is not a change in protein levels that prevents their identification in the *ppk* deletion strain, but rather, the absence of polyP itself. Intriguingly, one protein found enriched in WT lysates compared to WT lysates treated with yPPX, was the PPK enzyme itself. This association might serve a functional purpose, for instance, by promoting PPK-mediated conversion of ATP into polyP during stress and/or re-conversion of polyP into ATP ^35^ once nitrogen becomes available again. Incubation with RNaseA, however, only caused the depletion of a few proteins from the complex, suggesting that most proteins are scaffolded in the HMW complexes through polyP (Table S1). Subsequent gene ontology (GO) enrichment analysis of membership in the polyP-dependent Hfq complexes revealed a strong over-representation for proteins associated with ribonucleoprotein complex assembly, ribosome assembly, and RNA modifications (Fig. 4E). Of particular interest were 23 translation-associated proteins that showed a log2-fold enrichment in WT relative to *ppk* cells between +1.1 to +6.6 and included all three translation initiation factors (IFs), several ATP-dependent RNA helicases, numerous rRNA modifying enzymes and ribosomal subunits, as well as proteins related to RNA degradation (Table S1). Moreover, we also identified enrichments of chaperones, proteases, and select glycolytic proteins, including the RNA binding glyceraldehyde-3 phosphate dehydrogenase. Most intriguingly, however, we noted that the protein composition was highly reminiscent of the composition of eukaryotic stress granules and Processing (P)-bodies, which are equally enriched for proteins involved in RNP assembly, RNA stability and translation ^36^. Working at the functional level, we determined that 35% of the GO terms with which HP-body proteins are annotated overlap with those of P-body and/or stress granule proteins (Fig. 4E; Supplementary Dataset 1), reflecting significant enrichments relative to what would be expected by chance (p=0.004, 0.001, and 0004 for the P-body, stress granule, and joint overlaps, respectively, via resampling tests). Moreover, the set of terms observed to occur at the three-way intersection between these sets are particularly enriched for functionalities related to RNA metabolism, translation, and stress responses, all of which appear to be of great importance for the role of HP-bodies in promoting N-starvation. Importantly, the polyP-dependent enrichment of these proteins in HP-bodies could not be explained by differences in their expression levels according to our RNAseq expression analysis (Supplementary Dataset 2, “log2fc” tab); there is no detectable difference between the median levels of transcripts encoding HP-body proteins in the N24 condition relative to N0. The median increase in transcript levels under N24 starvation in WT vs. *ppk* cells is a log2 fold change of +0.26 (95% CI 0.15-0.37, p=3.63*10^-6^, Wilcoxon rank sum test), reflecting at most a modest stabilization of these transcripts (see below). Instead, our results suggest that the polyP-dependent recruitment of components of the translational machinery into Hfq foci is an active and specific process that likely contributes to the translational inhibition previously observed in N-starved bacteria^37^, and possibly analogous to the manner that eukaryotic P-bodies affect the stability of select RNAs. Based on these findings, we concluded that polyP has a profound impact on the protein composition of HP-bodies and we provide evidence for the role of HP-bodies as organizing centers for proteins involved in protein translation, RNA processing and decay functionally analogous to cytosolic condensates in mammalian cells.

**Figure 4.**
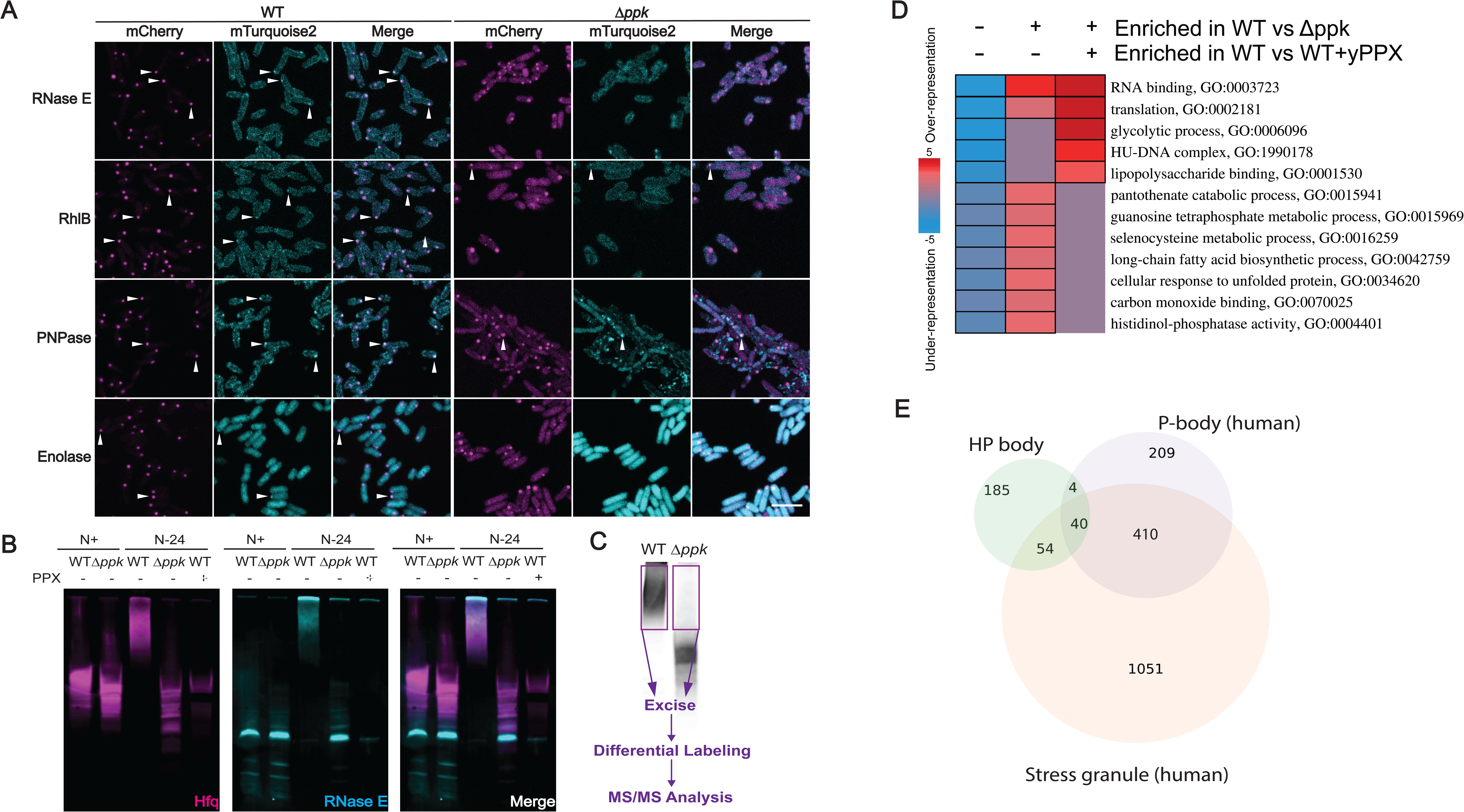
Protein composition of HP-bodies reminiscent of mammalian stress bodies. (**A**) Colocalization of Hfq-mCherry (magenta) and mTurquoise2-labeled RNA degradosome components (cyan) in N24 MG1655 *hfq::hfq-mCherry rne::rne-mTurquoise2*, *pnp::pnp-mTurquoise2*, *rhlB::rhlB-mTurquoise2* or *eno::eno-mTurquoise2* WT and Δ*ppk*. **(B)** Native western blot of N+ and N24 lysates from in MG1655 *hfq::hfq-mCherry rne::rne-mTurquoise2* WT relative to Δ*ppk* using antibodies against mCherry (Hfq) and GFP (RNase E). N24 samples were left untreated (-) or digested with PPX for 2 hours (+) prior to electrophoresis. **(C)** Diagram showing the workflow of MS/MS on native gel-resolved samples. Excised regions are indicated by purple rectangles. **(D)** Gene ontology terms of proteins enriched in WT samples relative to those of Δ*ppk* cells, in WT lysates relative to WT lysates treated with PPX, or both. Stronger red coloring indicates stronger enrichment in a given category. **(E)** Euler diagram showing the overlaps of GO terms annotated to proteins identified in the HP body vs. those in human P-bodies and stress granules (Supplementary Dataset 1).

### HP-bodies sequester and stabilize polyadenylated transcripts

Eukaryotic P-bodies sequester polyadenylated (polyA)-mRNAs in order to store and stabilize select mRNAs during stress conditions ^38^. In contrast to mammalian systems, however, where polyadenylation serves to stabilize mRNAs, polyadenylation in *E. coli* aids as a targeting signal for mRNA degradation by the degradosome, with the length of the poly-A tails positively correlating with transcript turnover ^39^. Previous work showed that Hfq interacts with polyA-mRNAs and helps to prevent their degradation by the RNA degradosome ^40^. To first determine whether polyA-transcripts are sequestered into HP-bodies analogously to P-bodies or remain associated with Hfq in the solute phase, we used fluorescence *in situ* hybridization (FISH). Indeed, we observed accumulation of polyA-transcripts in a number of HP-bodies in N-starved WT cells (Fig. 5A, arrowheads) whereas no such enrichment was observed in the absence of polyP. These results are consistent with Hfq’s general propensity to bind poly-A sequences, and strongly suggested that HP-bodies, like mammalian P-bodies, sequester polyA-mRNAs during N-starvation. To directly determine the effects of HP-body formation on the polyadenylation status of mRNAs during N-starvation, we next tracked the lengths of untemplated poly-A sequences present at the 3’ end of transcripts in both WT and *ppk* cells at the N0 and N24 conditions (see Methods for details). Although we observed a general decrease in mean poly-A tail lengths as cells entered deep N-starvation conditions, we noted an even more pronounced decrease in cells lacking polyP (Figure 5B). These results suggested that under N-starvation conditions, HP-body formation aids in stabilizing mRNAs with longer polyA tails, which would otherwise be degraded by the degradosome. Entirely consistent will all of our previous results, absence of *ppk* had no discernible systematic effect on the poly-A tail lengths at N0 (Fig. 5C). Of particular interest was also the finding that many of the genes with a particularly increased poly-A tail length in the WT cells compared to the *ppk* knockout cells encode for proteins involved in amino acid uptake and the translational machinery (Fig. 5D). These results offer a functional connection to the classes of proteins observed to be enriched in HP-bodies and suggest the possibility of a concerted regulation and stabilization of those genes by HP-body formation at both the transcript and protein levels.

**Figure 5.**
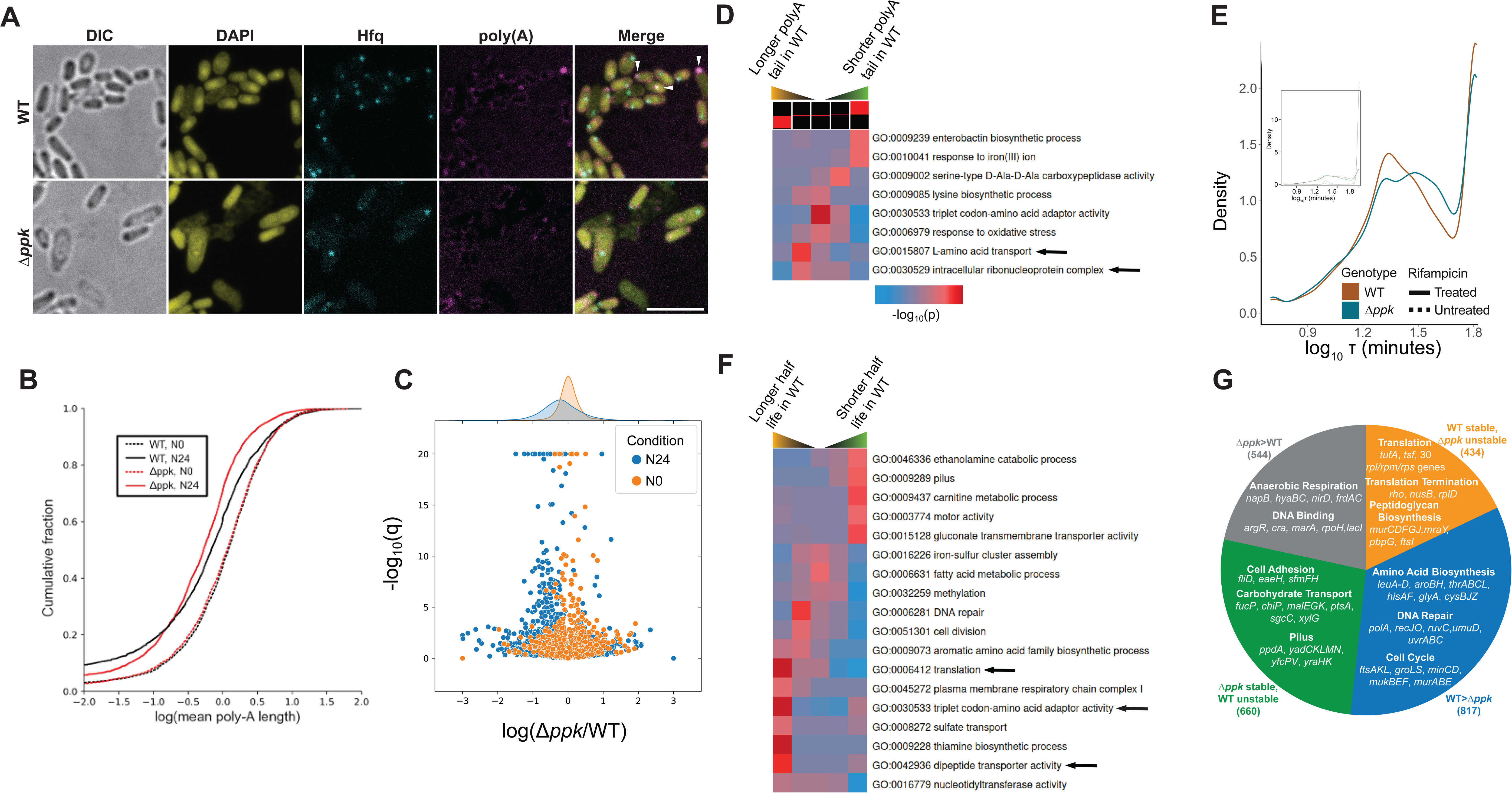
*Ppk* genotype affects global RNA polyadenylation and turnover. **(A)** Colocalization of poly-A RNA and Hfq as shown by fluorescent *in situ* hybridization against poly-A RNA in MG1655 *hfq::hfq-mCherry* WT and Δ*ppk* at N24. Scale bar indicates 5 µm. **(B)** Cumulative distributions of estimated mean untemplated poly-A lengths across the entire transcriptome in each of the indicated conditions. **(C)** Changes in poly-A tail length (x-axis) and statistical significance (y-axis) comparing *ppk* and WT cells under each of the indicated conditions; the leftward shift in *ppk* cells specifically at the N24 timepoint is clearly apparent. **(D)** Gene ontology (GO) term enrichments and depletions among the indicated quintiles of changes in polyA tail length upon deletion of *ppk*; terms in the lower quintiles show relative decreases in polyA tail length upon *ppk* deletion. **(E)** Distributions of the observed transcript half-lives in rifampicin treated MG1655 *hfq::hfq-mCherry* WT and Δ*ppk* (data in Supplementary Dataset 2). Decay constants outside of the range 5-65 minutes are clamped for plotting purposes. **Inset:** Comparison of untreated and treated distributions. **(F)** GO term enrichments and depletions among the indicated quintiles of changes in transcript half-life upon deletion of *ppk*. **(G)** Key GO terms and representative genes enriched in each of four major categories of *ppk*-induced stability changes (stable vs. unstable: τ− in one genotype > 65 min and < 55 min in the other; WT > *ppk* / *ppk* > WT: τ− in both genotypes < 65 min but τ− in one genotype increased by ≥10 min) along with representative genes (see Fig. S2C for full GO term analysis).

### PolyP regulates transcript stability by promoting HP-body formation

Based on these results, we concluded that the observed changes in poly-A tail lengths likely serve to differentially stabilize select mRNA transcripts during N-starvation, hence implicating HP-body formation in the regulation of mRNA stability. To directly monitor transcript stability, we added the transcriptional inhibitor rifampicin and conducted transcriptional shutdown experiments in N24 WT and *ppk* cells over a 60 min time course, using spike-ins for normalization^41^ (Fig. S2A, Supplementary Dataset 2). In both WT and *ppk* cells, we observed a clear bimodal distribution, with a split between apparently unstable transcripts (half-lives typically ranging from 10 to 50 minutes) and apparently stable transcripts (half-lives > 60 min, which we consider to be effectively non-decaying for the purpose of our analysis) (Fig. 5E; Supplementary Dataset 2). We calculated that about 28% of all transcripts identified were substantially less and about 27 % transcripts substantially more stable in the *ppk* mutant *versus* WT bacteria (Fig. S2B). Gene set enrichment analysis revealed several categories of genes for which transcript stability was systematically altered by deletion of *ppk* during N-starvation, including systematic stabilization of transcripts encoding genes involved in motility and surface adhesion, and broad destabilization of transcripts encoding genes involved in cell division, amino acid metabolism, and protein translation (Fig. 5F). Indeed, deeper analysis of specific genes of interest among the transcripts that are highly stable in WT during N-starvation but significantly destabilized in the *ppk* deletion strain showed enrichment for several components of the translational machinery (i.e., 30S small and large ribosomal subunits) as well as transcription elongation and termination factors (Fig. 5G and Fig. S2C). In tandem with our findings that the corresponding proteins are highly enriched in HP-bodies, we concluded that during N-starvation, polyP drives a systematic cellular response that sequesters the translational machinery and stabilizes key transcripts encoding this machinery. Consistently, transcripts that showed measurable turnover in both genotypes (exponential decay constant τ− < 65 min) but were still more stable in WT cells compared to the *ppk* deletion were enriched for cell cycle genes, chaperones, DNA repair and branched and aromatic amino acid biosynthesis (Fig. 5G). A completely different set of gene functions was represented among transcripts that were more stable in the absence of polyP and showed a high turnover in WT cells (Fig. S2C). These transcripts encoded for proteins involved in anaerobic metabolism, carbohydrate transport, and several extracellular adhesion factors. It is especially notable that the transcripts encoding HP-body components showed on average 44% larger increases in half-life (and thus higher stability) in the presence of polyP, relative to other transcripts (p=8.1*10^-5^, Wilcoxon rank sum test), reflecting a concerted preservation of the proteins involved at HP-bodies across all stages of synthesis and degradation. Taken together, these results strongly suggest that HP-body formation plays a major role in preserving transcripts that are necessary to rapidly restart protein synthesis and growth while targeting potentially less-essential transcripts for turnover. Finally, cross-referencing our data on transcript stability and polyA tail length, we found clear evidence that the transcripts involved in the core polyP-preserved processes such as translation are among the ones with increased polyA tail lengths and increased stability in WT bacteria compared to the *ppk* deletion mutant (compare Fig. 6D and F). These results strongly suggest that the polyP-dependent sequestration of the transcripts preserves the polyA tails and stabilized the transcripts during N-starvation.

**Figure 6:**
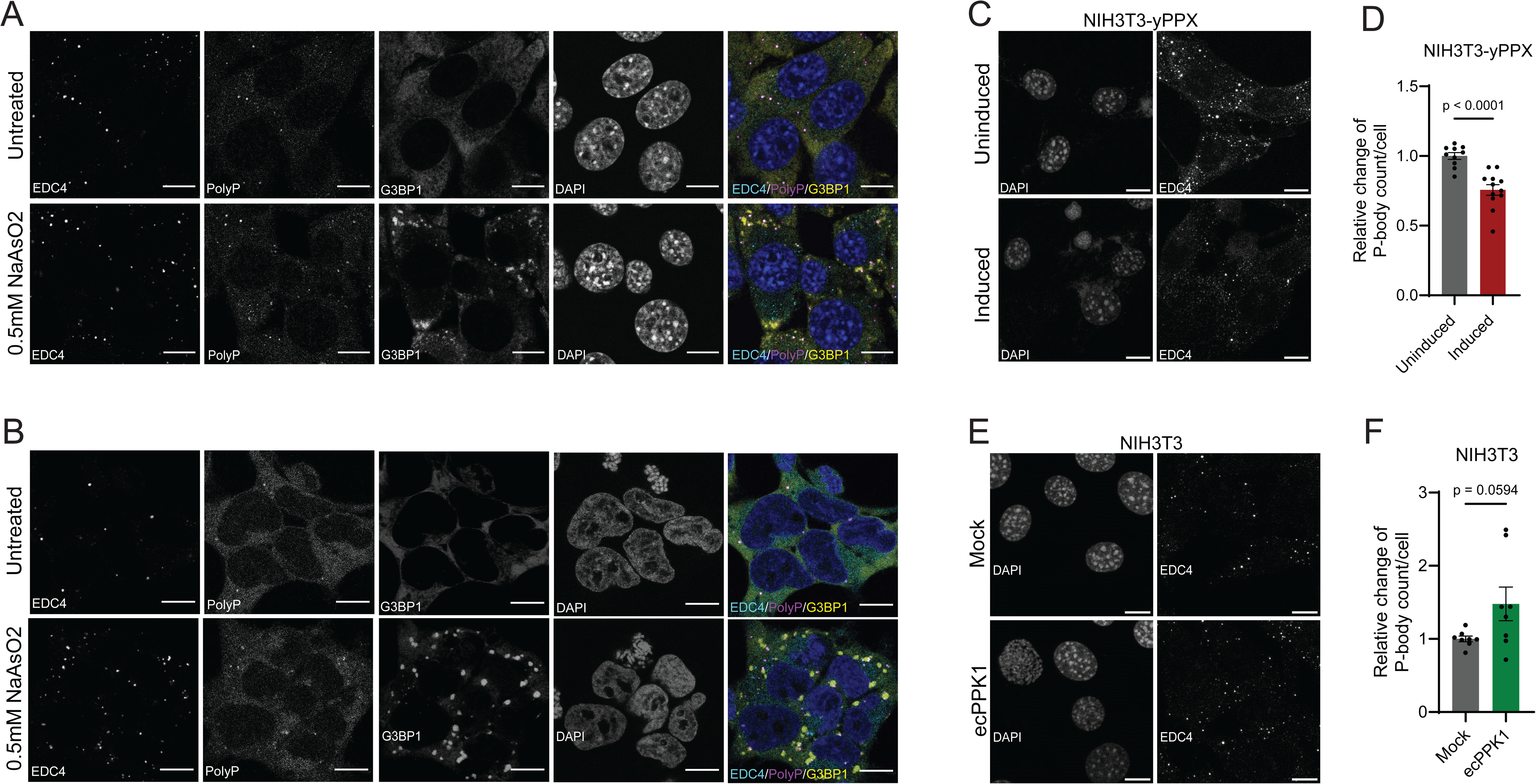
PolyP affects P-body formation in mammalian cells. (**A**) NIH3T3 and (**B**) HEK293 cells, either untreated or treated with 0.5 mM sodium arsenite (NaAsO_2_) for 30 min were fixed and stained for Processing (P)-bodies using antibodies against the marker protein EDC4 (cyan), stress granules (SG) using antibodies against the marker protein G3BP1 (yellow), polyP using the PPXBD-mcherry protein (magenta) and DAPI to visualize the nucleus (blue). A representative experiment is shown (n=3). (**C**) A stable NIH3T3 cell line expressing an inducible yeast exopolyphosphatase (yPPX) was fixed after 24h of addition of the inducer shield and stained for the P-body marker EDC4. (**D**) Quantification of the EDC4 positive punctate structures shown in (**C**). (**E**) NIH3T3 cells transiently transfected with the bacterial PPK1 was fixed after 24h of transfection and stained for the P-body marker EDC4. (**F**) Quantification of the EDC4 positive punctate structures shown in (**E**). Each data point represents the average P-body count per cell in an image of at least 20 cells. The average number of P-bodies counted in the uninduced (D) or untransfected (F) cells was normalized to 1. Image analysis was automated via thresholding to get an unbiased count of P-bodies. An unpaired t-test was used to assess statistical significance (n=3). All scale bars shown are 10 µm.

### PolyP plays a critical organizational role in eukaryotic P-body formation

Previous work revealed that in mammals, polyP is present in many different cell compartments, including the cytosol^26^. The compositional and functional overlap between HP-bodies and condensates in the cytosol of mammalian cells, particularly Processing (P-) bodies, which are involved in regulating RNA turnover, and stress granules (SGs) (Fig. 4E), which sequester the translational apparatus^42^, raised therefore the obvious question whether polyP is equally involved in eukaryotic condensate formation. We tested two cell lines, NIH3T3 (Fig. 6A) and HEK293 (Fig. 6B), which contain minor amounts of P-bodies even under non-stress conditions ^43,44^. Treatment with arsenite further increases their numbers and, at the same time, triggers the formation of the much larger stress granules ^45,46^. Fixation and IF-staining with antibodies against the P-body-specific marker protein EDC4^47^ and the SG-specific marker protein G3BP ^48^ aids in the visualization and differentiation of the two main cytosolic condensates. When we co-stained these cells before and after arsenite treatment using our mCherry tagged polyP-binding domain, we found a clear co-localization between polyP and EDC4 but not between polyP and the stress granule-specific marker protein G3BP1 (Fig. 6A, B). We did observe some clearly visible polyP-positive P-bodies in close vicinity to stress granules; however, these condensates have been observed before and suggested to be involved in exchanging select proteins and substrates ^49,50^. These results strongly suggested that polyP specifically interacts with components of P-bodies but not stress granules. To test whether manipulation of cellular polyP levels affects the formation of P-bodies, we constructed two NIH3T3-based cell lines; we generated stably transfected NIH3T3 cells expressing an inducible version of the yeast polyphosphatase yPPX, which significantly decreases cytosolic polyP levels within 24h of induction (Fig. S3A, B), or transiently transfected NIH3T3 cells with a vector encoding *E. coli* PPK, which significantly elevates the levels of polyP compared to non-transfected control cells (Fig. S3C, D). Indeed, and very similar to our results *in E. coli*, where depletion of polyP decreases and elevated levels of polyP increases HP-body formation, we observed a clear relationship between polyP levels and P-body formation in mammalian cells. Cells containing lower levels of polyP showed a significant drop in P-body numbers (Fig. 6C, D) whereas cells with higher-than-normal levels of polyP had increased P-body counts compared to non-transfected cells (Fig. 6E, F). These results suggest that polyP contributes to — and possibly even drives —P-body formation also in the context of mammalian cells, leading us to conclude that polyP’s role in condensate formation is likely to be evolutionary conserved.

## DISCUSSION

After more than a decade of extensive research, biomolecular condensate formation has been recognized as a critical subcellular organizing principle in both eukaryotes and prokaryotes. However, our understanding of these condensates is obscured by the lack of direct connections between their proposed cellular functions and observable phenotypes. Depleting cells of specific biomolecular condensates rarely causes detectable defects ^51,52^. Moreover, while many condensates are proposed to play a role in disorders such as cancer and neurodegenerative diseases ^53,54^, the vast majority of them fail to demonstrate a causal relation with the corresponding pathologies. In this study, we demonstrate a strong connection between the function of a condensate and its implications for cell survival. We show that HP-bodies, which form in response to nitrogen-starvation in *E. coli*, function as a central hub for RNA turnover, specifically sequestering and stabilizing transcripts involved in translation, the cell cycle and metabolism, while facilitating the degradation of RNAs less essential for survival and recovery (Fig. 7). Together with the sequestration of critical protein components of the translational and cell cycle machinery, HP-bodies provide bacteria with the ability to rapidly reinitiate protein synthesis and cell division after conditions become growth-permissive. This strong connection establishes HP-bodies as one of the few biomolecular condensates with clear physiological significance.

**Figure 7.**
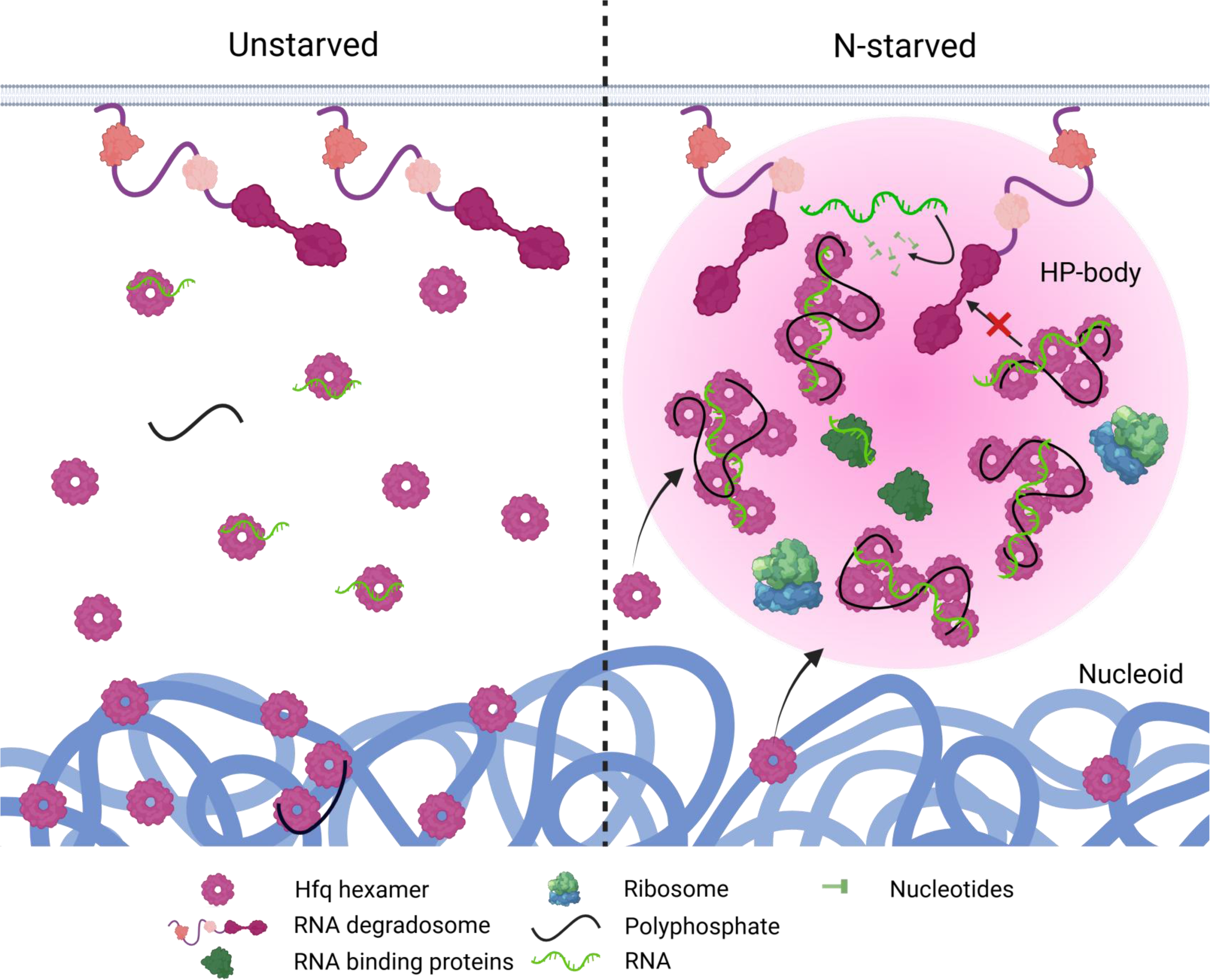
Current working model of HP-body formation and function. Upon N-starvation, *E. coli* accumulate long-chain polyP, which scaffolds Hfq-hexamers into higher oligomers, giving rise to phase separated condensates termed HP-bodies. PolyP promotes the selective sequestration of poly-adenylated RNAs and RNA-binding proteins into HP-bodies and facilities their association with the membrane-tethered RNA degradosome. HP body formation confers specific protection of transcripts involved in translation and metabolism while facilitating rapid turnover of transcripts that are likely less critical for survival.

Linear, negatively charged biopolymers such as RNA and DNA often participate in biomolecular condensate formation, where they mainly promote LLPS via multivalent electrostatic interactions with proteins ^23^. We now show that in bacterial cells, polyP — a prebiotic polyanion and likely the first high-energy compound on earth ^55^ — fulfils most (if not all) of the functions previously associated with nucleic acids in eukaryotic condensate formation. Long-chain polyP is necessary and sufficient to scaffold the RNA binding Hfq hexamers into higher-order oligomers, which increases the local concentration of Hfq and leads to Hfq condensation. By maintaining Hfq solubility, polyP prevents protein aggregation. On a functional level, we discovered that polyP binding affects several critical aspects of RNA metabolism, and that the HP-bodies accumulate at least some RNA species (notably those containing poly-A sequences) during nitrogen-starvation. Given that Hfq hexamers bind nucleic acids through several distinct interfaces, depending on the identity of the target nucleic acid, it is unsurprising that Hfq can simultaneously engage polyP and client nucleic acids, allowing the polyP to provide the multivalent interactions needed to drive phase separation while simultaneously engaging with some nucleic acid targets. It seems highly likely that differential affinity of polyP for the different nucleic acid binding regions of Hfq (rim, distal, proximal) might combine with competition for different subsets of nucleic acid targets in order to allow polyP to shape the condition-dependent landscape of Hfq-nucleic acid interactions. Further enumeration of the condition-dependent Hfq-RNA interactome, and the roles of different nucleic acid binding interfaces of Hfq in driving interactions with polyP and its competition with specific nucleic acid species, will be a promising area for future research.

Taken together, our observations establish polyP as a driving factor for HP-body formation and an essential regulator of subcellular organization in bacteria. Additionally, the compositional simplicity of polyP and the relatively small size of Hfq make HP-bodies a promising *in vitro* system for LLPS research. Our results showed that HP-bodies share many similarities with the cytosolic condensates found in many eukaryotes; like bacterial HP-bodies, mammalian P-bodies form around the SM-domain containing heptameric protein Lsm1-7, a eukaryotic counterpart of Hfq, accumulate mRNAs with short poly(A) tails and associate with components of the RNA degradation machinery to regulate turnover ^36,56^. Stress granules, on the other hand, share with HP-bodies the fact that they assemble specifically in response to stress conditions that trigger translation inhibition, and accumulate ribosomal subunits and translation initiation factors, indicative of stalled ribosomes ^47,57,58^. Although considered separate entities, P-bodies and SGs are often found in close spatial proximity and share select proteins and RNAs ^49^. Since HP-bodies combine features of both P-bodies and SGs on both compositional and functional levels, we propose that they might represent some of the most ancient forms of functional LLPS-driven condensates, and that polyP is the ancestral polyanion supporting phase separation. Given that polyP concentrations are highly sensitive to environmental conditions that lead to condensate formation, we now speculate that polyP is the quintessential and, based on its prebiotic origin, the ancestral scaffolding agent regulating condition-dependent formation of protein condensates *in vivo*.

## Resource Availability

### Lead Contact

Requests for further information and resources should be directed to and will be fulfilled by the lead contact, Ursula Jakob.

## Acknowledgments

We thank T. Shiba for providing the purified polyP of defined chain lengths and S. Wigneshweraraj for the strain containing PAmCherry-tagged Hfq. We thank K. Wan for purifying all the proteins used in this study, Kathrin Ulrich for her help with mass spectrometry sample preparation and Ellen Quarles for her help with the polyP quantification. We thank Hye Hyong Kweon for performing the mass spectrometry analyses. Usage of the Orbitrap Fusion Lumos instrument was funded by the Office of the Director, National Institutes of Health under Award Number S10OD021619. This work was supported by National Institutes of Health grant GM144731 (to J.S.B.), National Institutes of Health grant GM128637 (to L.F.), and National Institute of Health grant GM122506 (to U.J.).

## Author contributions

Conceptualization: J.G., L.F., and U.J. Methodology: J.G., R.L.H., J.S.B., L.F., and U.J. Investigation: J.R., R.L.H., A.R., C.A., and L.O.-R. Formal analysis: J.G., J.S.B., L.F., and U.J. Visualization: J.G., A.R., L.F., and U.J. Validation: J.G. and L.F. Writing— original draft: J.G. and U.J. Writing—review and editing: J.G., L.F., and U.J. Supervision: J.S.B., L.F., and U.J. Funding acquisition: J.S.B., L.F., and U.J. Project administration: U.J.

## Supplemental information titles and legends

**Figure S1:**
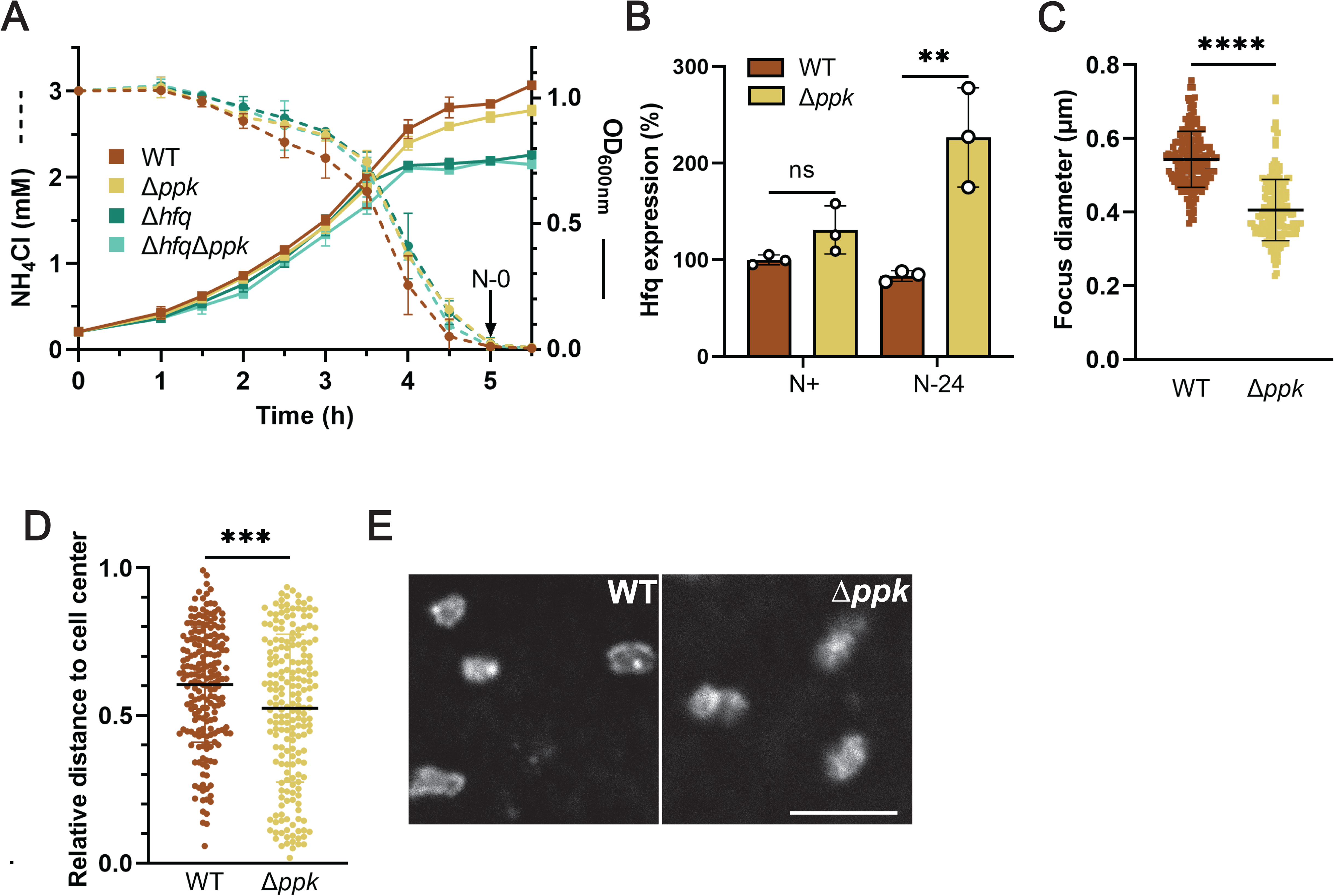
PolyP-mediated Hfq foc formation in *E. coli* (supplementing Figure 1) **(A)** Growth (solid lines) and nitrogen (N) levels (dashed lines) of WT, Δ*ppk*, Δ*hfq* and Δ*hfq*Δ*ppk E. coli* MG1655 in Gutnick minimal medium supplemented with 3 mM NH_4_Cl. Cells were grown at 37°C with aeration. Entry of N starvation (N0) is indicated. Error bars indicate SD (n=3). **(B)** Hfq-mCherry levels at N+ or N24 as determined by western blot using mCherry antibody. Total protein was used as loading control. Hfq-mCherry signal in N+ MG1655 *hfq::hfq-mCherry* cells was set to 100%. Error bars indicate SD (n=3). **(C)** Distribution of Hfq foci diameters in N24 in MG1655 *hfq::hfq-mCherry* WT and Δ*ppk* strains. Error bars indicate SD (n = 201). **(D)** Distribution of relative distances of Hfq condensates from 0 (cell center) to 1 (cell pole) in N24 in MG1655 *hfq::hfq-mCherry* WT and Δ*ppk* strains. Error bars indicate SD (n = 200). **(E)** Immunofluorescence images of fixed N24 WT or Δ*ppk E. coli* MG1655 *hfq::hfq-3xFLAG* using anti-FLAG antibodies for visualization. Comparisons in panels B-D were made by unpaired *t*-tests. ns, not significant; **p<0.01, ****p<0.0001, ****p<0.0001.

**Figure S2.**
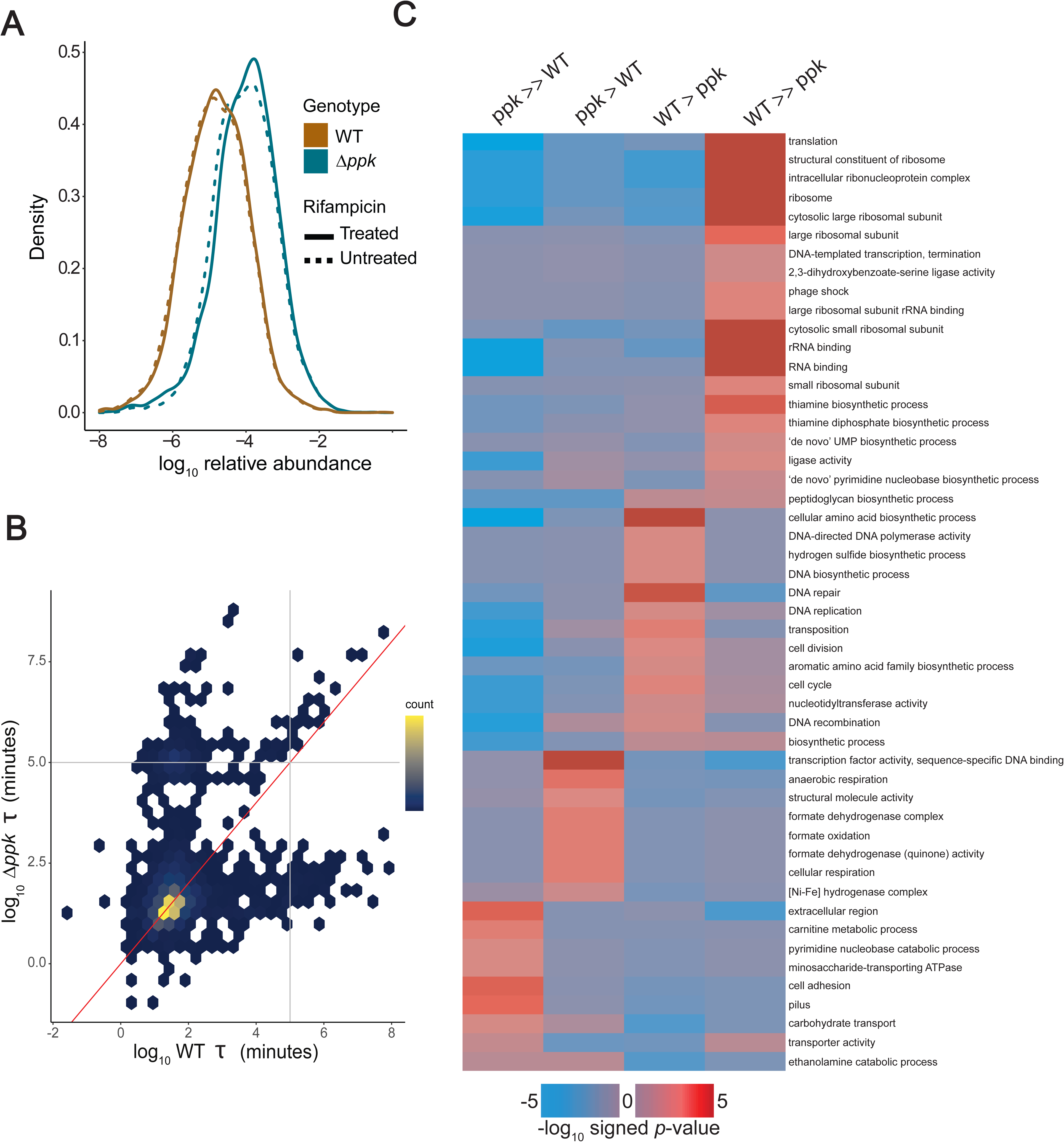
Effects of polyP status on transcript stability in the N24 stress condition (related to Figure 5). (**A**) Distributions of the observed transcript baseline abundances for the indicated combinations of genotype and rifampicin treatment. (**B**) Comparisons of fitted half-lives for transcripts in the WT vs *ppk* cell. (**C**) Full gene set enrichment analysis (performed using iPAGE software) on transcripts discretized into bins matching the categories in Fig. 5G.

**Figure S3:**
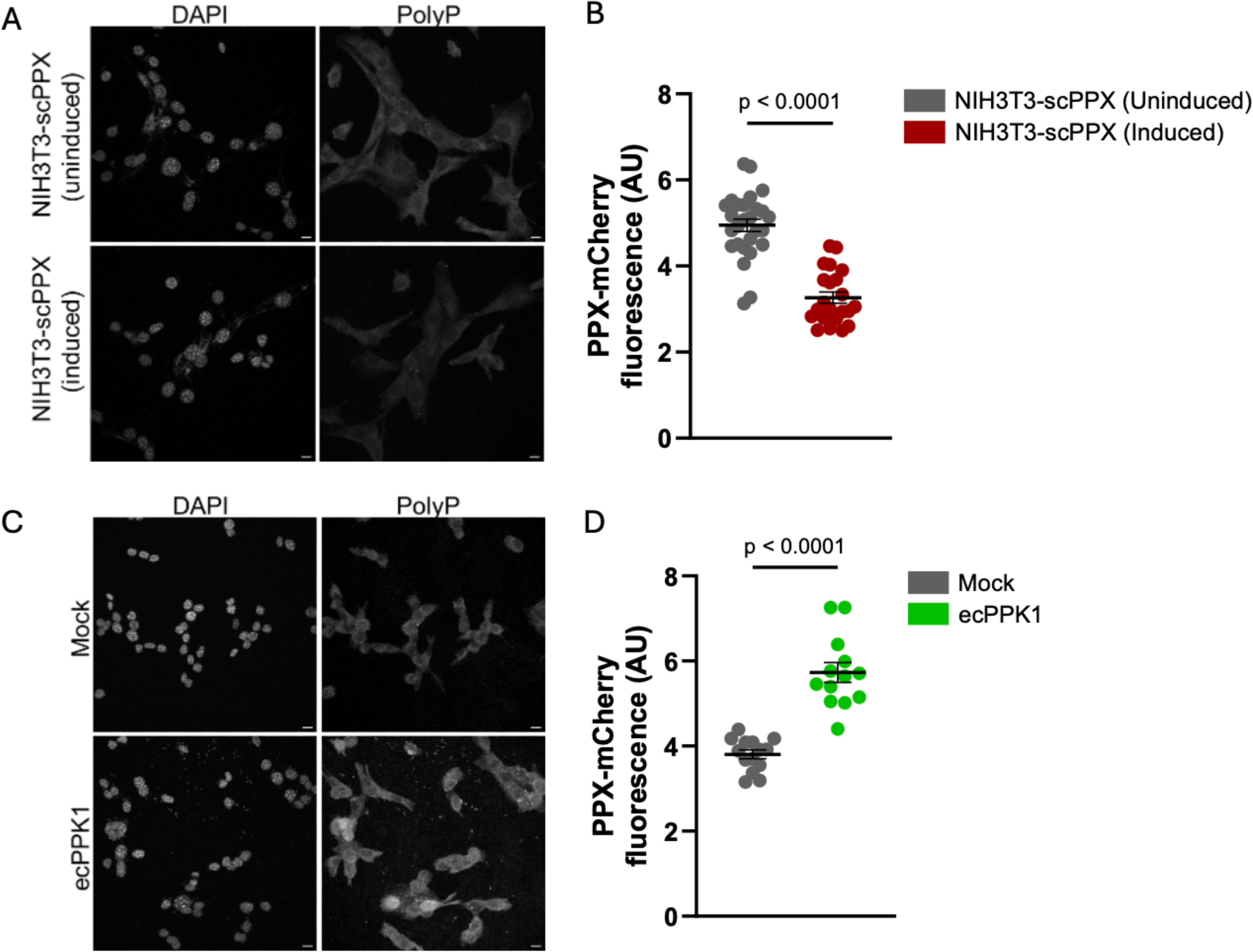
Manipulation of endogenous polyP levels in mammalian cells. (related to Figure 6). **(A)** A stable NIH3T3 cell line expressing a destabilization domain (DD) tagged yeast exopolyphosphatase (yPPX) was incubated in the absence (uninduced) and presence (induced) of small molecule, Shield1 (0.5 µM) that allows for the expression of yPPX. The cells were fixed and stained for polyP with PPXBD-mCherry. **(B)** PolyP levels were quantified based on fluorescence intensity of the PPXBD-mCherry probe. Each data point represents the average mCherry fluorescence intensity per cell (after background correction) in an image of at least 20 cells. **(C)** NIH3T3 cells were transiently transfected with a bacterial polyP kinase (ecPPK1) and were fixed and stained for polyP with PPXBD-mCherry. **(D)** PolyP levels were quantified based on fluorescence intensity of the PPXBD-mCherry probe. Each data point represents the mCherry fluorescence intensity of one cell (after background correction). An unpaired t-test was used to assess statistical significance (B, D). A representative set of images and quantification is shown (n=3). All scale bars are 10 µm.

**Table S1:**
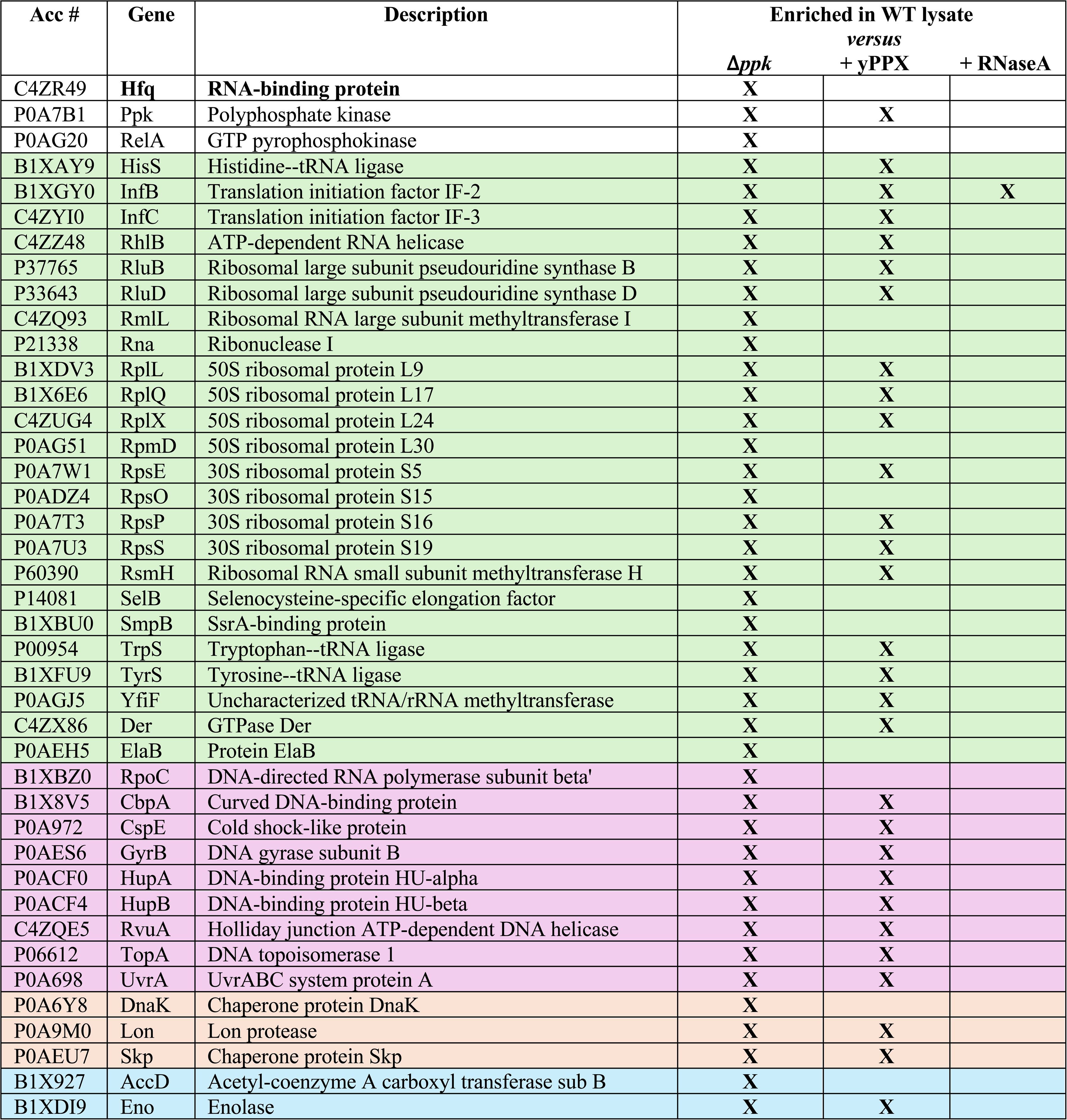
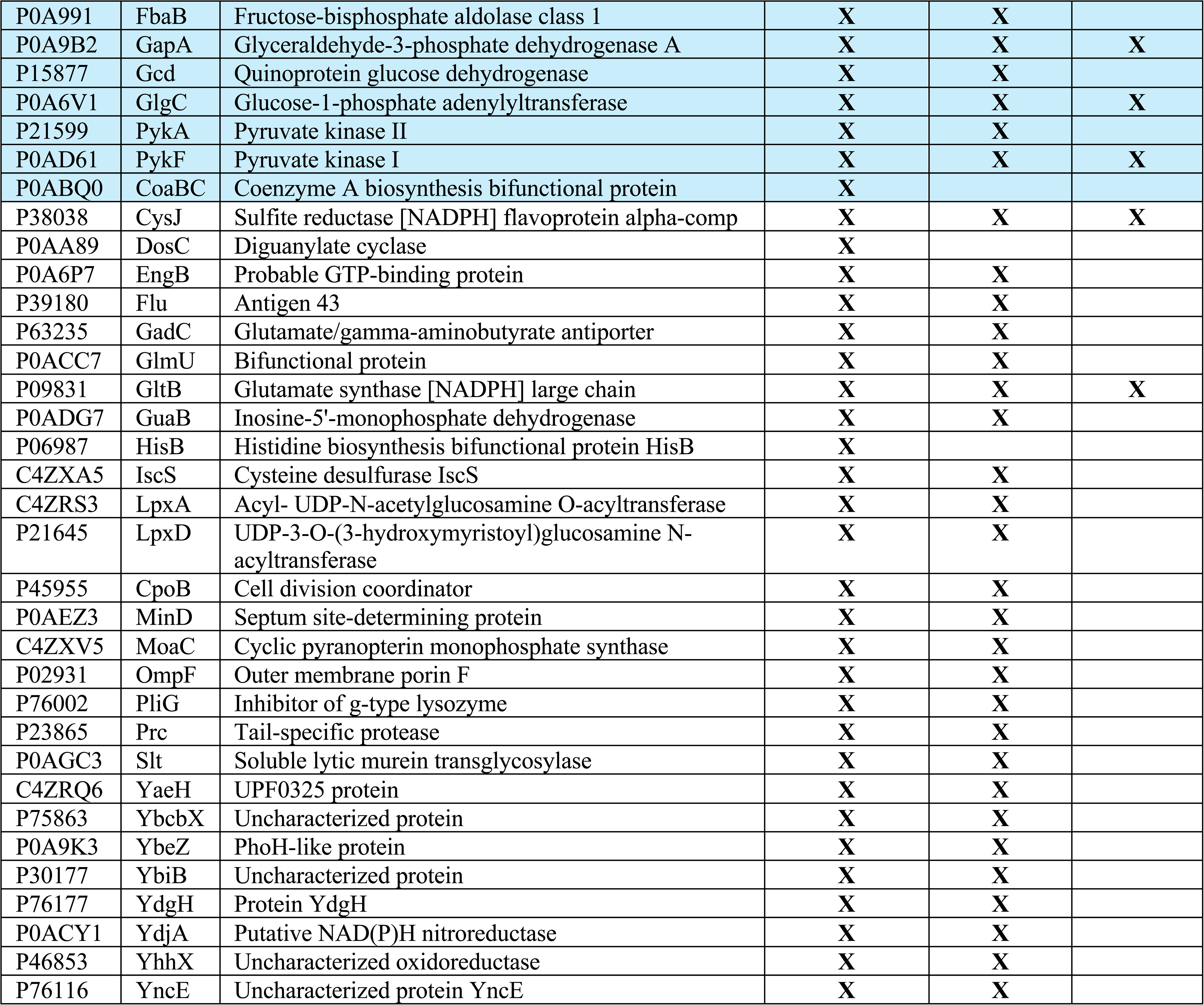
Proteins enriched in HP-HMW complexes (related to Figures 3, Supplementary data set 2). Lysates were prepared from N24 *hfq::hfq-*mCherry WT or the *hfq::hfq-*mCherry Δ*ppk* strain and analyzed on native PAGE. In addition, N24 *hfq::hfq-*mCherry WT lysate was treated with either yPPX to degrade polyP or RNaseA to degrade RNA prior to the native PAGE. Corresponding regions of the gels were excised, proteins pairwise differentially labelled (WT v *ppk*; WT v WT + yPPX; WT v WT + RNaseA) and analyzed by MS/MS analysis. Proteins with a log2 fold change (l2fc) >1 in at least 3 out of 5 replicates in WT v Δ*ppk*, 2 out of 4 replicates in WT v WT + yPPX or 2 out of 2 in WT v WT + RNase A are shown.

**Table S2:** GO-term enrichment analysis of HP body proteins (related to Fig, 4E). Shown are the set of GO terms present in the three-way interface between HP bodies, human P-bodies, and human stress granules.

|  |
| --- |
| **DNA-related** |
| GO:0003677 DNA binding |
| GO:0003697 single-stranded DNA binding |
| GO:0006281 DNA repair |
| GO:0006355 regulation of transcription, DNA-templated |
| **RNA-related** |
| GO:0003723 RNA binding |
| GO:0003724 RNA helicase activity |
| GO:0006401 RNA catabolic process |
| GO:0019843 rRNA binding |
| GO:0008298 intracellular mRNA localization |
| GO:1990904 ribonucleoprotein complex |
| GO:0008143 poly(A) binding |
| GO:0003676 nucleic acid binding |
| **Translation related** |
| GO:0003743 translation initiation factor activity |
| GO:0005840 ribosome |
| GO:0006412 translation |
| GO:0006417 regulation of translation |
| GO:0043022 ribosome binding |
| **Stress related** |
| GO:0009410 response to xenobiotic stimulus |
| GO:0009636 response to toxic substance |
| GO:0034605 cellular response to heat |
| GO:0042594 response to starvation |
| **Locations** |
| GO:0005576 extracellular region |
| GO:0005694 chromosome |
| GO:0005737 cytoplasm |
| GO:0005829 cytosol |
| GO:0016020 membrane |
| GO:0005886 plasma membrane |
| **Other** |
| GO:0000287 magnesium ion binding |
| GO:0005524 ATP binding |
| GO:0007165 signal transduction |
| GO:0008270 zinc ion binding |
| GO:0010468 regulation of gene expression |
| GO:0015031 protein transport |
| GO:0016301 kinase activity |
| GO:0016887 ATPase activity |
| GO:0042802 identical protein binding |
| GO:0042803 protein homodimerization activity |
| GO:0046872 metal ion binding |
| GO:0065003 protein-containing complex assembly |
| GO:0032991 protein-containing complex |

**Table S3.**
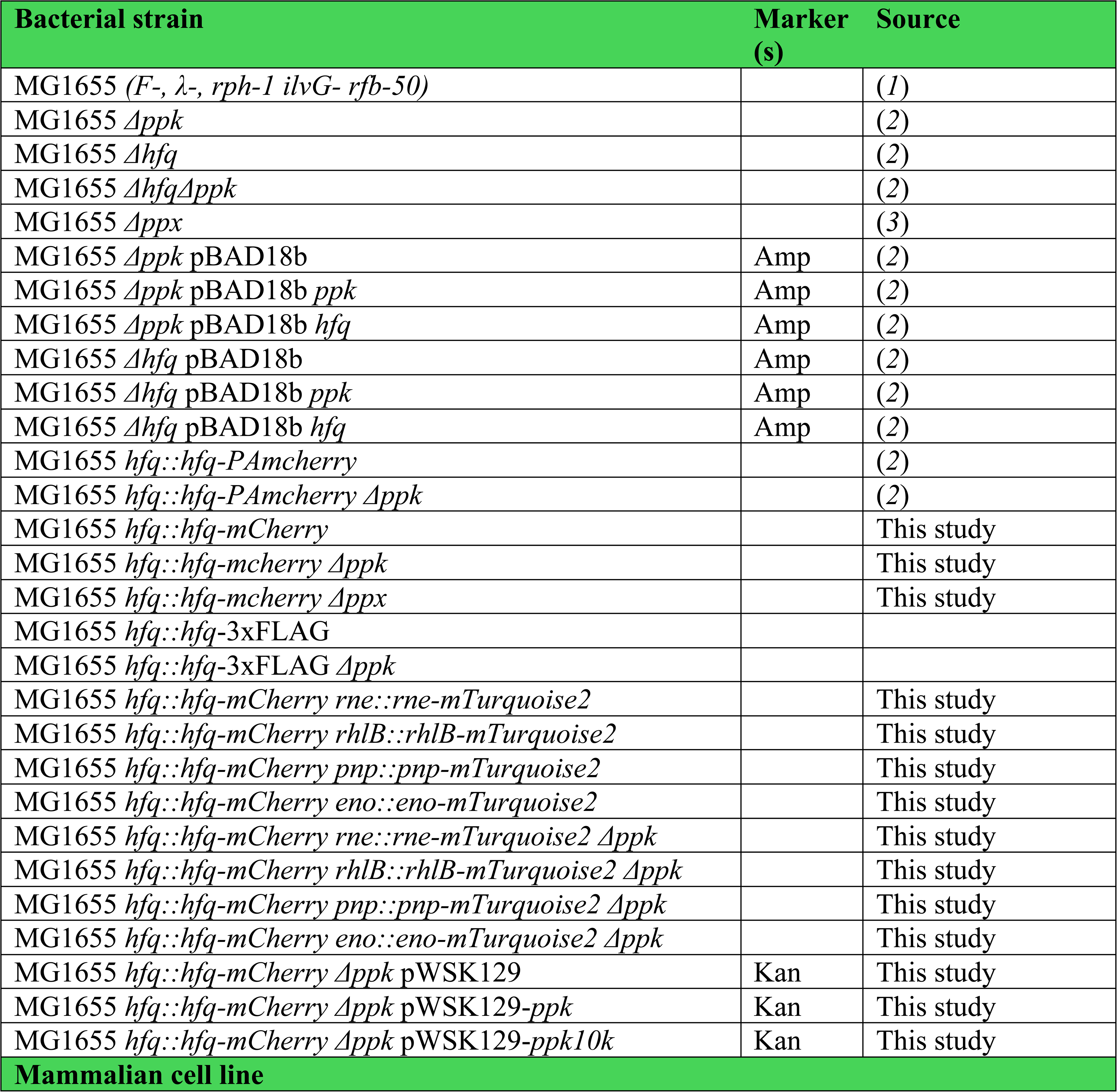
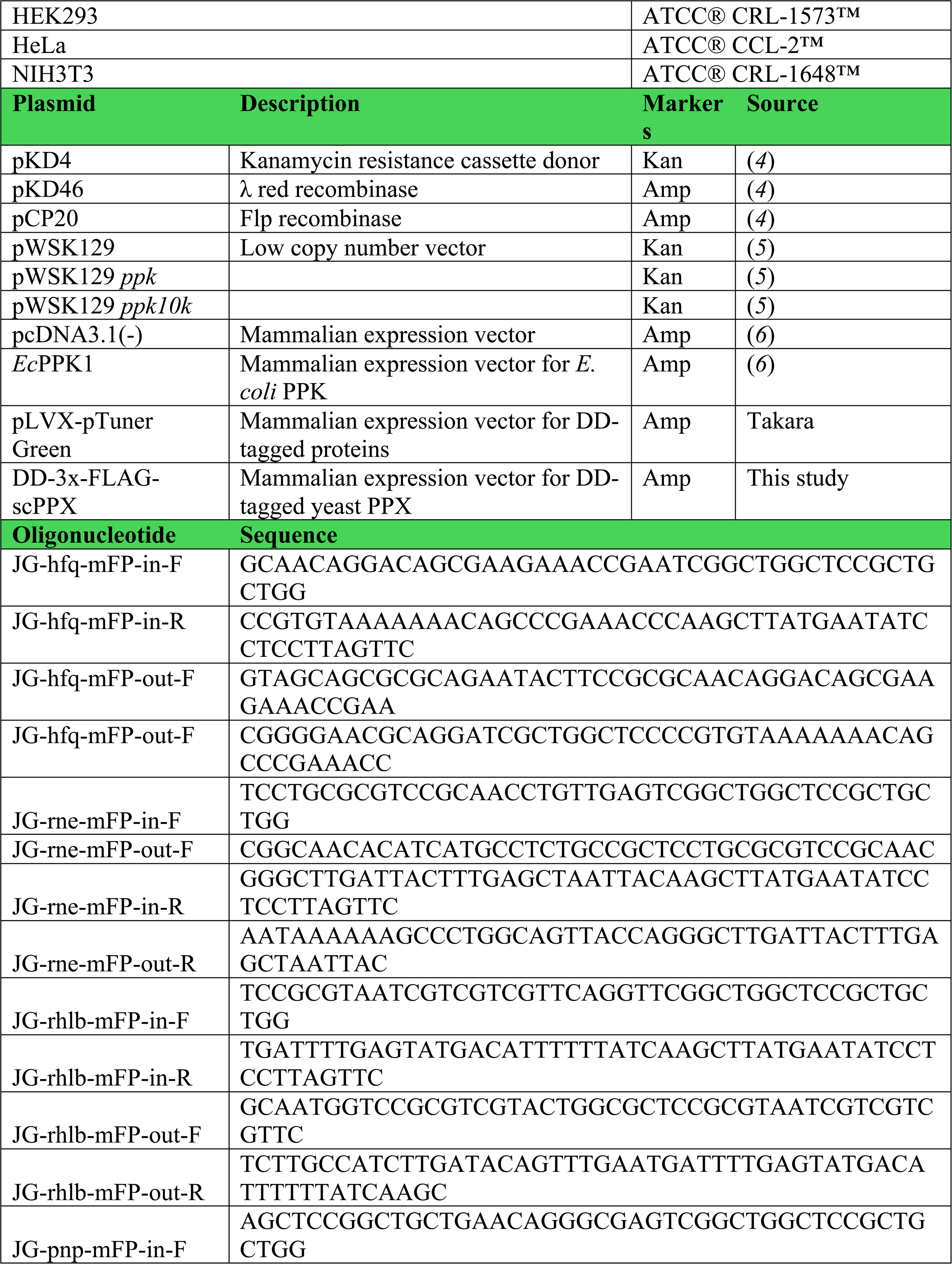
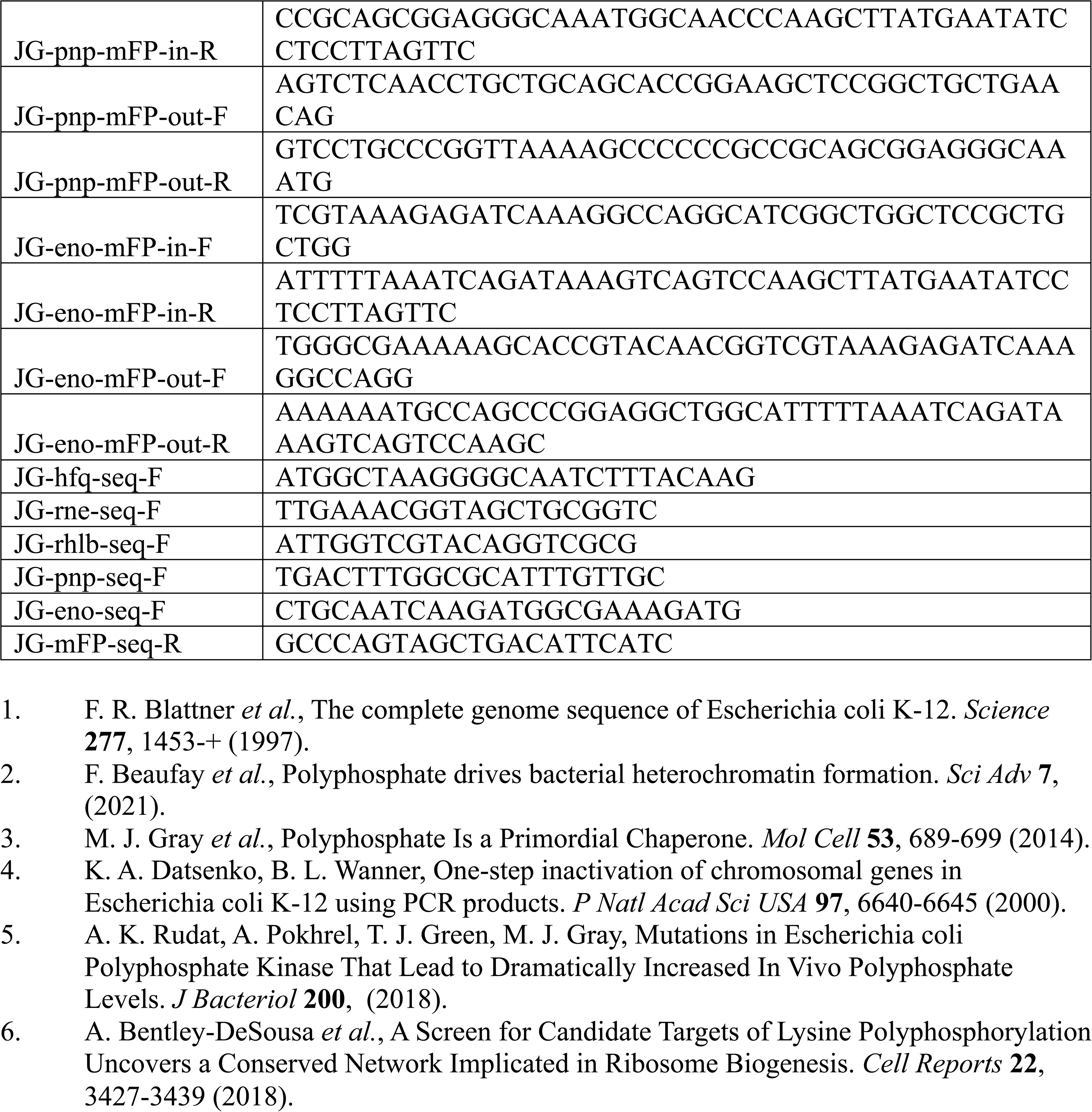
Strains, plasmids and cell lines used in this study.

**Supplementary Dataset 1:** Mass spectrometry results used for generating Table S1 and overlaps of GO terms annotated to proteins identified in the HP body vs. those in human P-bodies and stress granules (related to Fig. 4E). In the supplementary data file, the first tab gives more detailed descriptions of the data sets shown and the analysis used.

**Supplementary Dataset 2**: Fitted values and comparisons obtained from all high-throughput sequencing experiments used in this study. In the supplementary data file, the first tab gives more detailed descriptions of the data sets shown, the second tab gives the fitted values used in our analysis, and the third tab gives (specifically for the RNAseq datasets) q-values associated with each of the log fold changes given in the second tab.

## STAR Methods

### Bacterial strains and plasmids

All strains, plasmids and oligonucleotides used in this study were derived from *E. coli* strain MG1655 and are listed in Table S3. Alleles were deleted or fluorescently labelled via λ Red recombination method ^59^. Labelled alleles were transduced into various mutant strains using P1*vir* bacteriophages from appropriate donors. All plasmids used in this study were constructed with restriction digestion and ligation or Gibson assembly and verified by sequencing.

### Bacterial growth conditions

WT and mutant bacteria were grown in Gutnick minimal medium (33.8 mM KH_2_PO_4_, 77.5 mM K_2_HPO_4_, 5.74 mM K_2_SO_4_, 0.41 mM MgSO_4_) containing 0.4% (w/v) glucose, 10 mM NH_4_Cl and M9 trace elements (N+ medium) ^60^. For N-starvation experiments, overnight cultures in N+ medium were inoculated in Gutnick minimal medium containing 0.4% (w/v) glucose, 3 mM NH_4_Cl and M9 trace elements. For osmotic stress experiments, cultures at mid-exponential phase in LB medium were spun down, resuspended in equal volume of LB medium containing 1.17 M NaCl, and let grow for 4 hours. Under all conditions, bacteria were grown at 37°C with 200 rpm shaking. During growth in N-medium, the ammonium level in the medium was determined by using the Ammonia/ammonium microplate assay kit following the manufacturer’s protocol (#orb545637, Biobyrt). When indicated, kanamycin (100 µg/ml), ampicillin (200 µg/ml) and/or chloramphenicol (50 µg/ml) were added. Optical density of the culture was measured on a UV spectrophotometer. The number of viable cells in the culture were measured as CFU/ml after serial dilutions on LB agar plates and incubation overnight at 37°C.

### Fluorescence microscopy and FRAP measurements

Imaging of fluorescently labelled bacteria was performed on a Leica SP8 inverted microscope with 100× oil immersion objective, driven by LAS X software (Leica GmbH, Mannheim, Germany). Bacteria were grown under the conditions described above. At the indicated time points, 1 mL of cells were spun down, washed and resuspended in a small volume of Gutnick minimal medium without glucose or NH_4_Cl. Cells were loaded onto a cover slip and immobilized on 1% agarose pad made from the same medium ^12^. For imaging of N+ cells, medium with 3 mM NH_4_Cl and 0.4% w/v glucose was used for pad preparation. For imaging of the nucleoid, cells were stained with 1 µg/mL 4’,6-diamidino-2-phenylindole (DAPI, #D1306, Thermo Fisher Scientific) before being spun down and washed. For quantification of the foci, a minimum of 100 cells per sample was analyzed. All foci quantification and size measurements were performed with ImageJ. FRAP measurements were performed with the Zoom In mode, using the 405 nm and 587 nm lasers together at 100% intensity. Three pre-bleach images were acquired, followed by photobleaching and imaging every 5 sec for 115 sec. Fluorescence intensity was measured by LAS X software. Recovery data were plotted and fitted against a one-phase association curve with GraphPad Prism.

### Bacterial immunofluorescence microscopy in fixed cells

N24 cultures of WT and Δ*ppk hfq::hfq-mCherry* strains were cultivated as described. Cells equivalent to 1 ml OD_600_=1.5 were collected and fixed with fresh 4% paraformaldehyde (#1578100, Electron Microscopy Sciences) for 15 min at room temperature followed by 30 min on ice. Cells were spun down and washed three times in phosphate buffered saline (PBS), resuspended in 1 ml GTE buffer (50 mM glucose, 10 mM EDTA pH 8.0, 20 mM Tris-HCl pH 7.5) containing 20 µg/ml lysozyme, and incubated for 1 min. 20 µl of cells were loaded onto a poly-L-lysine (#P8920, Sigma-Aldrich) coated coverslip (#72230-01, Electron Microscopy Sciences) for 15 min. Coverslips were washed three times with PBS, and dried at room temperature. After rehydrating with PBS, samples were blocked with 2% bovine serum albumin (#A3059, Sigma-Aldrich) in PBS for 20 min. A PPXBD-GFP probe ^33^ was diluted in blocking solution to 10 µg/mL, and samples were incubated at room temperature overnight. After five washes in PBS, IRDye 680RD-labeled goat anti-mouse antibody (#926-32210, LI-COR Biosciences) was added in a 1:1,000 dilution and incubated for 2 hours at room temperature. Coverslips were washed three times in PBS, sealed on a glass slide with a drop of antifade reagent (#9071S, Cell Signalling) and imaged as previously described.

### Single-molecule fluorescence microscopy

WT and Δ*ppk* Hfq-PAmCherry strains colonies were cultured in LB medium for 6 hours at 37°C and 200 rpm. Subsequently, the cells were diluted 1:400 into N+Gutnick and grown overnight under the same conditions. To induce nitrogen-starvation, the overnight culture was diluted in N-Gutnick to an OD_600_ ∼0.02 and grown as above. Cells were imaged at 6- and 32-hours post N-Gutnick inoculation. For imaging preparation, 0.5 ml of cells were centrifuged and resuspended in filtered spent N-Gutnick media. The cells were immobilized on 2% (w/v) agarose:spent N-Gutnick medium pads and imaged at room temperature on a wide-field Olympus IX71 inverted microscope equipped with a 100x 1.40 NA oil immersion objective. Hfq-PAmCherry molecules were photoactivated with 50 to 100-ms pulses of 405-nm laser (Coherent Cube, 405-100; 2 W cm^-2^, followed by imaging under excitation by a 561-nm laser (Coherent Sapphire, 561-50; 0.34 kWcm^-2^). Frames were captured at 50 Hz using a 512 × 512-pixel Photometrics Evolve EMCCD camera. The recorded movies were subsequently analyzed using the SMALL-LABS algorithm ^61^ for single molecule localization and tracking. To identify H-body foci, all frames for each movie were summed to generate a composite image. The foci present in this image were detected by the Laplacian of Gaussian algorithm (*min_sigma*=3, *max_sigma*=5, *threshold* = 0.1), segmented and assumed to be H-bodies. Putative foci were filtered to determine if indeed these are stable H-bodies. A focus was kept if it contained at least ten trajectories of at least ten frames each where at least 70% of the trajectory overlapped with the focus. The median trajectory length in our dataset is approximately ten frames; therefore, this number was chosen for the minimum threshold of trajectory length in the filtering. To identify the different states of Hfq, single-molecule trajectories were then classified based on their overlap with foci areas. Trajectories were classified as ‘In’ if they completely overlapped with a focus area for their entire duration, ‘In/out’ if they overlapped with a focus for 25% to 99% of their duration, and ‘Out’ if they overlapped for less than 25% of their duration^62^. The squared displacement, *r^2^*, was calculated for consecutive localizations within a trajectory of at least 4 frames (*τ−* = 20 ms). The cumulative probability of the squared displacements in the observation period *τ−* (*P*(*r^2^, τ−*)) was generated from the pool of squared displacements across multiple tracks for each sample replicate by counting the number of squared displacements less than or equal to *r^2^* normalized by the sample size. The CDF of *r^2^* was fit to analytical functions describing the diffusive processes with three dynamics states. *D_1_*, *D_2_*, and *D_3_* are the diffusion coefficients for the different states, and *α* describes the relative fraction between the states.

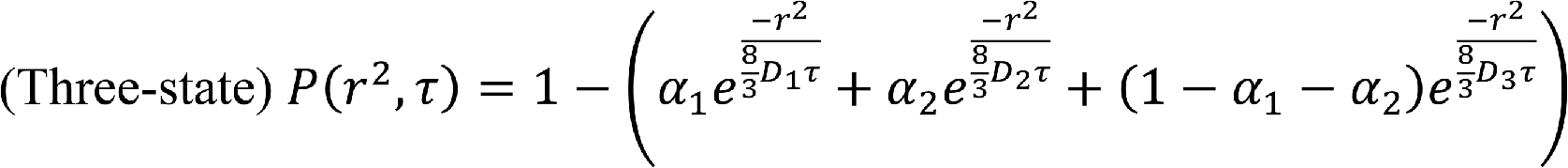

Here, we assumed the *D_3_* population corresponds to freely diffuse molecules as *D_3_* > 1 μm^2^/s, such that the fraction of freely diffuse molecules is *F_free_* = *α*_3_. We assumed the two slower populations, *D_1_* and *D_2_*, correspond to a combination of nucleoid-associated and within-condensate Hfq molecules. To identify the specific contributions from within-condensate molecules, we first analyzed the trajectories that were classified as “In”. The same diffusion analysis was performed as above but using a two-state model.

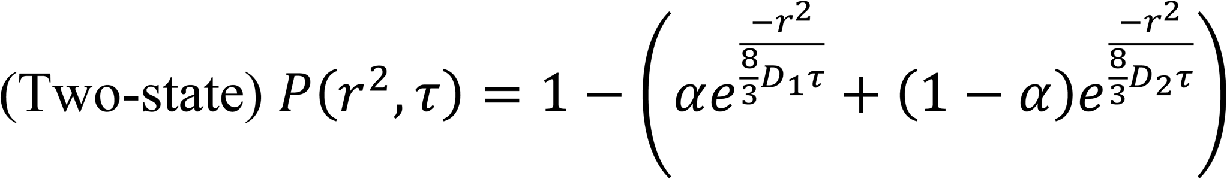

The resulting *D_1_* and *D_2_* were consistent with the *D_1_* and *D_2_* from the entire trajectory dataset. Next we fit the combined “In” and “In/out” subsets to a three-state model and used the combined weight fractions *α_1_ + α_2_* as the within condensate contribution for this subset as *D_3_* was consistent with freely diffuse molecules. The total fraction of Hfq in condensates is then defined as:

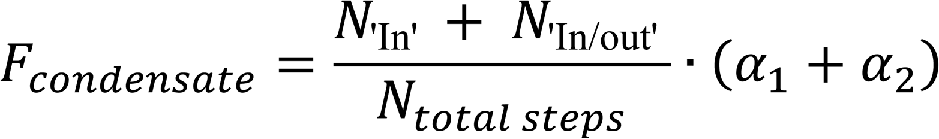

Since it is possible for a condensate focus to overlap with the nucleoid, we cannot exclude that some amount of *F_condensates_* is also nucleoid-associated. We defined the fraction of Hfq that is only nucleoid-associated and not within a condensate as:

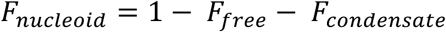

### PolyP extraction and detection

PolyP was extracted as previously described ^63^ with modifications. Briefly, WT and *Δppk* bacteria cultured overnight in N+ medium was reinoculated into N-medium and grown at 37 °C. At exponential phase (OD_600_=0.5), N0, N-3, N6, N-12, N-18 and N24, cultures equivalent to 4 mL OD_600_=1.0 were collected, spun down, washed in PBS and resuspended in 350 µL AE buffer (50 mM sodium acetate, 10 mM EDTA). Suspension was sonicated on ice, and 300 µL phenol and 40 µL 10% SDS was added. The mixture was incubated at 65 °C for 5 min, and then on ice for 2 min. 300 µL chloroform was then added, and the mixture was spun down. The upper aqueous phase was mixed with 350 µL chloroform, spun down, and aqueous phase was again collected. After measuring sample volume, 3 times volume of 100% ethanol, 10% volume of 3 M sodium acetate and 5 µL 5mg/mL glycogen (#AM9510, Thermo Fisher) was added. The mixture was vortexed and incubated on ice for 1 hr. DNA, polyP and glycogen was pelleted, washed with 70% ethanol and air dried. Dried samples were dissolved in 500 µL MP buffer (125 mM NaCl, 5 mM MgCl_2_, 25 mM Tris-HCl pH 7.5). Per 50 µl sample, 1 µl M-SAN nuclease (#70950-202, ArcticZymes) was added and incubated at 37 °C for 2 hr. To examine polyP on gel, 10 µL sample was mixed with 2 µL 20% glycerol and 2 µL gel loading dye, loaded onto a 4-20% TBE gel (#EC6225Box, Invitrogen), and ran for 4 hr at 100V. PolyP 300mer and 130mer were loaded as chain length markers. To measure polyP concentration (in phosphate units), 30 µL M-SAN digested samples were added with 1 µl *Saccharomyces cerevisiae* exopolyphosphatase (yPPX) and incubated at 37 °C for 2 hr. Potassium phosphate solutions ranging from 0 to 250 µM was used as standards. In a microplate, 25 µL of each standard was mixed with 25 µl MP buffer, 15 µl 5x yPPX buffer (100 mM Tris-HCl pH 7.5, 25 mM MgCl_2_, 250 mM ammonium acetate), 10 µL water and yPPX. 25 µL of each sample was mixed with 50 µL water. Molybdate working solution was prepared by mixing 91.2% total volume of detection base (0.6 mM antimony potassium tartrate, 600 mM H_2_SO_4_, 2.4 mM ammonium heptamolybdate) and 8.8% total volume of 1 M ascorbic acid. 25 µL working solution was added to each standard and sample well, rested for 2-5 min, and absorbance at 882 nm was measured.

### Immunoblotting

Strains of interest were grown to various stages of N-starvation, and cultures equivalent to 1 ml OD_600_=0.6 were collected, spun down, washed and resuspended in 50 µL SDS-polyacrylamide gel loading buffer, and lysed by incubating at 95 °C for 10 min. Protein samples were then run on a 4-12% NuPAGE Bis-Tris gel (#NP0336BOX, Thermo Fisher) at 175V for 45 min. The gel was then transferred onto a polyvinylidene difluoride membrane (#1620174, Bio-Rad) and blocked with 5% milk in TBST for ≥1 hour. The membrane was incubated in a dilution of 1:1000 anti-mCherry monoclonal antibody (#M11217, Thermo Fisher) or 1:1000 anti-GFP polyclonal antibody (#632592, Takara Bio) in blocking solution for ≥1 hour, rinsed five times with TBST, and then incubated in 1:10000 diluted anti-rat or anti-rabbit secondary antibodies labeled with IRDye 680RD or 800CW (#926-68076 and #925-32211, LI-COR Biosciences). The membrane was rinsed three times in TBST, once in TBS, and imaged on a LI-COR Odyssey CLx imager (LI-COR Biosciences). All washes and incubations were done at room temperature. A gel was run in parallel and stained with Coomassie blue as normalization control. Image quantification was performed using ImageJ, and three biological replicates were included in each experiment.

### Native western blot

Bacterial cultures equivalent to 40 ml OD_600_=1.0 were collected, spun down, and lysed by sonication in 5 mL native lysis buffer [20 mM Tris-HCl pH 8, 1 mM β-mercaptoethanol, 1 mM EDTA, 2% glycerol and cOmplete protease inhibitor (#5056489001, Sigma-Aldrich)]. Samples were centrifuged to remove insoluble debris, and supernatant was mock treated or digested with DNase I (#18068015, Invitrogen), RNase A (#10109169001, Sigma-Aldrich) or yPPX for 2 hours at 37°C before running on a 4-20% TBE gel (#EC6225Box, Invitrogen) at 100V for 16 hours. The gel was incubated in SDS-PAGE running buffer for 20 min and was subjected to western blotting following standard protocols described above.

### Fractionation of soluble and insoluble proteins

As previously described ^64^, cultures equivalent to 4 ml OD_600_=1.0 were collected by centrifugation. The pellet was resuspended in 50 µL ice-cold lysis buffer (10 mM potassium phosphate pH=6.5, 1 mM EDTA, 20% sucrose, 1 mg/mL lysozyme, 50 u/ml benzonase), incubated for 30 min on ice, and frozen at -80 °C. After thawing on ice, 360 µL ice-cold buffer A (10 mM potassium phosphate pH=6.5, 1 mM EDTA) was added. The mixed sample was transferred to a 2 mL tube containing 200 µL 0.5 mm glass beads and was incubated for 30 min at 8 °C with 1,400 rpm shaking. After settling down the beads, 200 µl lysate was centrifuged at 16,000×g for 20 min, and supernatant (soluble proteins) was collected for further use. The pellet (insoluble proteins) was washed with buffer A, buffer B (buffer A containing 2% Nonidet P-40), and again with buffer A. Previously collected supernatant was mixed with ¼ volume of 100% trichloroacetic acid, incubated for 10 min at 4 °C and spun down at 21,000×g, and precipitates were washed three times with ice-cold acetone. After washing acetone was removed from samples by heating at 37 °C. Both soluble and insoluble proteins were resuspended in 100 µL 1× reducing SDS buffer (6.5 mM Tris-HCl pH=7.0, 10% glycerol, 2% SDS, 0.05% bromophenol blue and 2.5% β-mercaptoethanol), boiled at 95 °C for 10 min, and wash subjected to further analysis as described above.

### RNA-seq

WT and Δ*ppk* Hfq-mCherry strains were streaked onto LB agar plates from cryogenic storage and grown at 37°C. Colonies (three biological replicates per strain) were inoculated into N+ Gutnick medium and grown overnight at 37°C with shaking at 200 rpm. The overnight culture was back-diluted to OD_600_=0.01 in 250 mL N-Gutnick medium. At 0, 6, 12 and 24 hours after N-starvation, cultures equivalent to 40 mL OD_600_=1.0 were collected and immediately spun down for 3 min at 13,000×g at 4°C in a fixed-angle rotor. The cell pellets were shock-frozen in a dry ice-ethanol bath, and stored at -80°C. Immunoprecipitations were conducted following a protocol adapted from ^21^. Three biological replicates of samples were extracted in three separate experiments (one per replicate). Each pellet (∼ 40 O.D. * mL) was resuspended in 600 µL of lysis buffer (10 mM Tris pH 8.0, 1x cOmplete™, EDTA-free Protease Inhibitor Cocktail (Roche, cat. No. 11836170001), 50 mM NaCl, 3 µL of Ready Lyse lysozyme per mL) and incubated for 15 minutes at 30°C. Samples were placed on ice, then sonicated in the ice bath at 25% amplitude for 20 seconds (5 seconds on, 15 seconds off). The samples were clarified by centrifugation (16,000 r.c.f.,10 minutes, 4°C), and the liquid transferred into new microfuge tubes. To each sample, an equal volume of 2x IP buffer (200 mM Tris, pH 8.0, 600 mM NaCl, 4% (v/v) Triton X-100 with 2× protease inhibitors (Roche, cat. No. 11836170001) and 0.2 mg/ml BSA) was added and mixed by inversion. To preclear samples, 100 μL of pre-washed (add 100ul of beads per sample in microfuge tube, place on magnetic stand, remove storage buffer, add 1 mL 1X IP buffer, mix by inversion, clear using magnetic stand, remove liquid, resuspend in 100 µL of 1x IP buffer per sample) protein G magnetic beads (NEB; lot 101325116) were added to each sample and incubated for 2 hours on a nutating platform at 4°C. Beads were cleared using a magnetic stand, and the liquid was transferred to new 2 mL microfuge tubes (the preclearing step is included to ensure compatibility with a set of immunoprecipitation experiments that will be reported separately). 100 µL of each sample was transferred to a new tube containing 400 µL of ChIP elution buffer (50 mM Tris, pH 7.0, 10 mM EDTA, 1% (w/v) SDS), mixed, and kept on ice until proteinase K (10 µl per sample; Fermentas, 50 mg/ml) digestion. After mixing by inversion, all tubes were incubated for 2 hr at 37°C. This was followed by phenol/chloroform extraction cleanup (500 μL of acidic 5:1 phenol/chloroform (VWR, E277) was added, mixed, and allowed to separate into layers at 16,000 r.c.f., 5 minutes, 4°C); the water layer was transferred to a new tube with 500 μL of 24:1 chloroform/isoamyl alcohol, mixed, allowed to separate as before at 16,000 r.c.f., 5 minutes, 4°C); finally, the water layer was transferred to a new tube). Nucleic acids from the samples were precipitated by the addition of 1/25 volume of 5 M NaCl, 2 μL of Glycoblue coprecipitant (Invitrogen), 1 volume of 100% (v/v) ethanol and 1 volume of 100% (v/v) isopropanol, mixed, and then incubated for 1 hour at 4°C, then > 2 hr (up to overnight) at - 20°C. Nucleic acids were pelleted via centrifugation (16,000 r.c.f., 15 minutes, 4°C), washed once with ice-cold 95% (v/v) ethanol, and air dried. Pellets were resuspended in 83 μL of 0.1X TE (1 mM Tris pH 7.0, 0.1 mM EDTA) and stored at -80°C. The DNA was removed from the samples 10 μl of 10X Baseline-ZERO Buffer plus 5 μL of Baseline-ZERO DNase (Lucigen) and 2 μl of murine RNase Inhibitor (NEB), and incubated 37°C for 30 min. The RNA was collected/cleaned using the Zymo RNA Clean and Concentrate Kit-96 per kit directions (Zymo Research) and eluted with either 15 (extracts) or 20 μL (inputs) of water. The concentration of RNA was determined with 3 μL of each sample using QuantiFluor RNA System (Promega) and the provided standards.

### Library preparation for RNA-Seq

The material described above was subsequently treated with NEBNext rRNA Depletion Kit (Bacteria) [NEB]. For each input, 11 µl of diluted material (0.15-0.44 µg) was used. Depletion was performed per kit directions except that Zymo RNA Clean and Concentrate Kit-96 [Zymo Research, per kit directions (3000 r.c.f., 5 min centrifugation) and eluted with 11 μL of water] was substituted for the post DNase I digestion bead clean-up in an attempt to preserve small RNAs that may be present. Library preparation was performed using NEBNext Ultra II Directional RNA Library Prep Kit for Illumina (NEB) and 5 μL of rRNA depleted. Samples were prepared per kit directions except that samples were fragmented for 1 min, the post-second strand synthesis bead clean-up was replaced with an Oligo Clean and Concentrate kit [Zymo Research, per kit directions (3000 r.c.f., 5 min centrifugation) and eluted with 25 μl of water, and add 25 μl of water prior to end prep reaction], diluted (1:100) NEBNext Unique Dual Index UMI Adaptors DNA Set 1 were used instead of the NEBNext Adaptor for Illumina, the post ligation bead binding involved 174 μl of Axyprep beads (Axygen) and 68 μl of isopropanol, and the NEBNext Primer Mix was used for the PCR amplification step. Pooled libraries were subjected to Illumina sequencing on a NextSeq platform with 38 x 37 bp paired end reads.

### RNAseq Data Analysis

Raw RNA-seq reads were pre-processed using the IPOD-HR pipeline version 2.7.1 ^70^, acting through the preprocessing and alignment stages. In brief, this includes adapter removal, quality score trimming alignment using bowtie2, and quality control using FastQC plus additional custom metrics. After preprocessing and alignment, overlaps of aligned reads to genes were quantified using the summarizeOverlaps() function of the R package GenomicAlignments (version 1.26.0) running under R 4.0.2, with options “singleEnd = FALSE, inter.feature = FALSE, ignore.strand = TRUE, mode = “IntersectionStrict”,fragments = TRUE”, with the MG1655 transcriptome defined according to the Operon Set from RegulonDB (downloaded March 2023). Differential expression calling was then performed using DeSeq2 ^71^, with a model including terms for the sample genotype, timepoint (0h, 6h, 12h, or 24hr), sequencing preparation batch, interactions between genotype and timepoint. Multiple hypothesis testing correction was performed using the IHW method ^72^.

### Rifampicin treatment, sequencing, and data analysis for RNA stability measurements

#### Growth and harvest

N24 WT and Δ*ppk E. coli* were supplemented with 150 µg/mL rifampicin (#R3501, Sigma-Aldrich) and incubated at 37°C with 200 rpm shaking. Samples were collected before rifampicin addition, or at 10, 30 and 60 min after treatment, and were immediately mixed with RNAProtect^®^ Bacteria Reagent (#1018380, Qiagen); for the post-treatment timepoints, a separate untreated control was collected after the same lag time. Cells were pelleted and stored at -80°C. Each sample was also plated for CFUs at the time of harvest in order to provide a normalization factor (used below). Three biological replicates were performed for each genotype/condition combination.

#### Lysis and RNA purification

On ice, each sample pellet was resuspended in 90 µl of 1X TE (10 mM Tris pH 7.0, 1 mM EDTA pH 8.0), briefly centrifuged to remove liquid from tube walls and transferred to 1.5 ml tube and then transferred to 1.5 ml microfuge tubes. Then 1 µl of Ready-Lyse Lysozyme Solution (LGC Biosearch Technologies) was added to each sample and mixed. Samples were incubated at 30°C for 15 min and returned to ice. Then, 10 µl of Proteinase K (Thermo-Fisher) was added to each sample and mixed. Samples were incubated at 23°C for 15 min, with resuspension every two minutes then returned to ice. ERCC RNA spike-in control mix (Ambion) was diluted 1:25 in water as one batch (8.6 µl of ERCC, 206.4 µl water). Then, 5 µl of the diluted mix was added to each sample, except for *Δppk* t=60 (no rifampicin) replicate 1, which received 1.6 µl instead of 5 µl. After vortexing the samples, they were clarified by centrifugation (12,000 r.c.f., 30 sec.) and up to 250 µl of clarified lysate was transferred to a new tube. The samples were cleaned using RNA Clean & Concentrator-96 (Zymo Research) per manufacturer’s directions for total RNA clean-up and eluted with 25 µl of water per sample. Each sample was transferred to tubes that contained 60 µl of water, 10 µl of 10X Baseline Zero DNase buffer, 5 µl of Baseline Zero DNase (LGC Biosearch Technologies), and 2 µl of RNase Inhibitor, Murine (recombinant, NEB) in each tube. Samples were incubated at 37°C for 30 min. The samples were re-cleaned using RNA Clean & Concentrator-96 (Zymo Research) per manufacturer’s directions for total RNA clean-up and eluted with 20 µl water per sample. The concentration of RNA was determined using 5 µl samples and Quantifluor RNA system (Promega) per manufacturer’s directions for quantitating RNA in multiwell plates.

#### Library preparation and sequencing

Appropriate amounts of nuclease-free water were added to 0.1 µg of RNA per sample to achieve 11 µl of starting material for NEBNext rRNA Depletion Kit (Bacterial, NEB). The kit was used per manufacturer’s directions to remove rRNA from the samples. Library preparation was performed using NEBNext Ultra II Directional RNA Library Prep Kit for Illumina (NEB). Samples were prepared per kit directions except that 2.5 µl of diluted (1:100) NEBNext Unique Dual Index UMI Adaptors DNA Set 1 was used instead of the NEBNext Adaptor for Illumina, and that Omega Mag-Bind TotalPure NGS Beads were used instead of NEBNext Sample beads for post second-strand synthesis, post adapter ligation, and post PCR clean-up steps. Pooled libraries were subjected to sequencing on a NextSeq 2000 instrument.

#### Data analysis

After initial read preprocessing, alignment and QC using the IPOD-HR processing pipeline (equivalent to that described above for the RNA-seq samples), overlaps of aligned reads to genes were quantified using the summarizeOverlaps() function of the R package GenomicAlignments (version 1.26.0) running under R 4.0.2, with options “singleEnd = FALSE, inter.feature = FALSE, ignore.strand = TRUE, mode = “IntersectionStrict”,fragments = TRUE”. A composite reference transcriptome including both all annotated MG1655 transcripts and the transcripts from the ERCC spike-in set was used. Read counts for each feature (gen) were then normalized by the total read count for that sample, and then sequencing depths of genomic features further rescaled by the median abundance of ERCC transcripts in that sample (considering only ERCC transcripts with a relative abundance greater than 5*10^-4^) in order to provide a relative estimate of per-cell transcript abundance. For each gene, we then fitted an exponential decay equation (*y*=*a*\*exp(-1\**t*/*b*), where *y* is the normalized transcript abundance, *a* and *b* are coefficients to be fitted, and *t* is the time) to the data for each time course, using the scipy.optimize.curve_fit function from scipy 1.8.0, using the ‘dogbox’ optimizer with both parameters constrained to be non-negative, and *a* and *b* initialized to 0.001 and 20, respectively; *a* represents a baseline abundance (at t=0) for each transcript, and *b* is the exponential decay time constant (in minutes). We considered ‘measurable’ decay constants to be those between 5 minutes and 65 minutes.

### Untemplated poly-A tail length determination

WT and Δ*ppk* Hfq-mCherry strains were streaked on LB agar plates from cryogenic storage and grown at 37 °C. Colonies (four biological replicates per strain) were inoculated into N+ Gutnick medium and grown overnight at 37 °C with shaking at 200 rpm. Overnight culture was back-diluted to OD600=0.01 in 250 mL N-Gutnick medium. At OD600=0.4∼0.45 (N+) and 24 hours after nitrogen-starvation (N-24), cultures equivalent to 40 ml OD600=1.0 were collected and immediately spun down for 3 min at 13,000×g at 4 °C in a fixed-angle rotor. Cell pellets were shock-frozen in a dry ice-ethanol bath, and stored at -80 °C.

#### Sequencing Library Preparation

##### RNA extraction and cleaning

To lyse cells, 700 µl of DNA/RNA Shield (Zymo Research, R1100-50) was added to each pellet, cells were resuspended, incubated at room temperature for 5 min, and then frozen solid at -80C. Upon thawing, lysate was transferred to 2 mL and clarified via centrifugation (12000 r.c.f, 5 min). The soluble fraction was transferred to new tubes. Samples were cleaned using RNA Clean & Concentrator-96 (Zymo Research) per manufacturer’s directions for RNA clean-up from samples in DNA/RNA Shield. The cleaned samples were digested with BaselineZero DNase (1X Baseline Zero buffer, 10 µL of Baseline Zero enzyme, 2 µL of Murine RNase Inhibitor [NEB, M0314]) for 30 min at 37°C. Samples were cleaned using an RNA Clean & Concentrator-96 (Zymo Research) per manufacturer’s directions for total RNA clean-up. The cleaned samples were digested a second time with BaselineZero DNase (1X Baseline Zero buffer, 10 µL Baseline Zero enzyme, 2 µL of Murine RNase Inhibitor (NEB, M0314)) for 60 min at 37°C. RNA was cleaned by the addition of 1.8X volumes of RNAClean XP beads (Beckman Coulter), mixing, incubating on ice for 15 min, removing the unbound fraction, washing undisturbed beads with 200 µL of 80% ethanol twice, removing the ethanol and allowing evaporation of the residual liquid, followed by elution of RNA in 25 µL of water. Samples were stored at -80°C.

##### RNA ligation and conversion to dsDNA

For each sample, 1 µg of RNA and water to reach a 40 µL total volume were added to a PCR strip tube. The 5’ phosphates were removed from the RNA by rSAP (1X rCutsmart buffer, 3 units of rSAP, [NEB, M0371]; 80 units of murine RNase Inhibitor [NEB, M0314]) incubated for 2 hr at 37°C. RNA was cleaned by the addition of 1.8X volumes of RNAClean XP beads (Beckman Coulter), mixing, incubating on ice for 15 min, removing the unbound fraction, washing undisturbed beads with 200 µL of 80% ethanol twice, removing the ethanol and allowing evaporation of the residual liquid, followed by elution of RNA in 7 µL of water. The RNA was transferred to new PCR strip tubes. T4 RNA ligase I (30 units, NEB, M0437M) was used to ligate a 5’ phosphorylated, 3’-dideoxy terminated, UMI containing adaptor oligo (100 pmol of 5Phos/TCAAGCAGTAGNNNNNNNNCAGCAGTTCGATAAGCGG/3ddC, synthesized by IDT) to the 3’ ends of the RNA in a buffered reaction (1X T4 RNA Ligase Reaction Buffer, 12.5 % PEG 8000, 1 mM ATP, 40 units of murine RNase Inhibitor, 1/10 volume DMSO) at 16°C for 12-14 hr. The ligation product was cleaned by the addition of 1.8X volumes of RNAClean XP beads, mixing, incubating on ice for 15 min, removing the unbound fraction, washing undisturbed beads with 200 µL of 80% ethanol twice, removing the ethanol and allowing evaporation of the residual liquid, followed by elution of RNA-DNA in 7 µL of water. The RNA-DNA eluate was then transferred to new PCR strip tubes. The RNA of the RNA-DNA was fragmented in buffer (4 µL of 5x Protoscript II buffer, M0368S, NEB; 1 µL of 10mM dNTPs) with a primer complementary to the adaptor oligo (5’CCGCTTATCGAACTGCTG, 20 pmol per reaction) at 94°C for 6 min prior to reverse transcription. The RNA-DNA was reverse transcribed using ProtoScript II Reverse Transcriptase (200 units of M0368S, NEB) which was mixed into each reaction after the addition of DTT (10mM final) and murine RNase Inhibitor (8 units) to each sample. The reactions were incubated 25°C for 5 min, 42°C for 1 hr, 80°C for 15 minutes, and then 4°C. After samples were placed on ice, 8 µL of NEBNext Second Strand Synthesis Reaction Buffer with dUTP (E7426AA, NEB), 1 µL of random primers (E7422AA, NEB), and 47 µL of water were mixed well into each 20 µL RT reaction. Then 4 µL of NEBNext Second Strand Synthesis Enzyme (E7425AA, NEB) was mixed into each reaction. Reactions were incubated at 16°C for 1 hr. The dsDNA was cleaned by the addition of 1.8X volumes of Mag-Bind TotalPure NGS beads (Omega Biotek), mixing, incubating 5 min RT, removing the unbound fraction, washing undisturbed beads with 200 µL of 80% ethanol twice, removing the ethanol and allowing evaporation of the residual liquid, followed by elution of the DNA in 50 µL of water.

##### Library Preparation and sequencing

The DNA was prepared for sequencing using NEBNext Ultra II DNA Library Prep Kit for Illumina (E7645L, NEB) with NEBNext Multiplex Oligos for Illumina Dual Index Primers Set 1 (E7600S, NEB) per manufacturer’s protocol (with a 1 to 10 dilution of the standard adapter) except the blue capped USER enzyme (E7428AA, NEB) was used instead of red capped USER enzyme (E7602AVIAL, NEB), Mag-Bind TotalPure NGS beads (Omega Biotek) were used instead of NEBNext Sample Purification Beads, and 14 cycles of amplification were performed at the PCR stage. Pooled reactions were sequenced using NextSeq 1000 Sequencing System (Illumina) using a P2 100 cycle kit with 75 bp read 1, 25 bp read 2, 19 bp index 1, and 8 bp index 2 reads.

#### Data Analysis

Reads were initially pre-processed and aligned using the same procedures as those noted above for the standard RNA-seq datasets, with the crucial exception that reads were aligned in local, rather than end-to-end, mode. Apart from this, the number of ‘T’ bases in the read (which, due to the location/orientation of our sequencing adapter, correspond to 3’-end ‘A’ bases in the original RNA) was taken to be the number of terminal As on that read. For each gene, we then aggregated the distribution of 3’-terminal ‘A’ counts occurring on reads mapping (in the correct orientation) to any position from 50 bp before to 250 bp after the end of that gene, thus assigning reads occurring in that 3’ window to the gene that they are close to the end of. We note that while we could instead have performed this analysis at the level of annotated transcripts/transcriptional units, it appeared likely to us that in many cases we might miss polyA tail assignments due to imperfect knowledge/enumeration of all transcripts produced under the conditions being studied, whereas the primary weakness with our current approach is that we might assign to a single gene a polyA state that also affects other genes in the same operon. For each gene, we then fitted zero-inflated negative binomial regression models to the distributions of aggregated polyA tail length observations across the 16 experiments performed here (four biological replicates for each of four conditions). We incorporated coefficients contributing to the mean polyA tail length for the possible combinations of *ppk* genotype and nitrogen-starvation status (four degrees of freedom) and fitted only a single zero inflation rate and overdispersion parameter for each gene. Fits were performed using the python statsmodels package, version 0.13.2, with default parameters except that we allowed a maximum of 500 iterations for coefficient fitting.

### Motif Analysis

All motif analysis was performed using FIRE 2.0 software ^73^ using default settings except as otherwise noted. For continuous inputs (e.g., log2 fold changes), inputs were split into five equally populated bins for the mutual information analysis; for discrete inputs, bins were allocated as shown. In addition to bin-specific p-values, FIRE provides strength metrics for motifs based both on motif z-scores and robustness in a resampling test; filtering based on either or both criteria are described with each motif enrichment application.

### GO Term Enrichment Analysis

All GO term enrichment analysis was performed using the iPAGE 1.2b package ^74^ using default settings except as otherwise noted. The GO term annotation set used was derived from the Uniprot ^75^ MG1655 reference proteome indexed by b-number for any nucleic acid based analysis, and for all Uniprot proteins associated with *E. coli* K12 (taxon ID 83333) indexed by Uniprot identifier for the mass spectrometry analysis. For continuous inputs (e.g., log2 fold changes), inputs were split into equally populated bins for the mutual information analysis; for discrete inputs, bins were allocated as shown.

### Mass spectrometric (MS) analysis

For MS analysis of native gel, samples were collected and lysed in the same manner as native western blot. Two 4-20% TBE gels (#EC6225Box, Invitrogen) were run in parallel at 100V for 1.5 hours. One gel was subjected to native western blotting to identify the location of Hfq-mCherry oligomers, while the other gel is stained in warmed Coomassie blue for 5 min, destained overnight, and washed with ddH_2_O for 24 hours. Area corresponding to the Hfq-mCherry oligomers was cut out from the WT N24 lane, while areas of the same migration distance on other lanes were also collected. For analysis of co-IP samples, samples were run on a 4-12% NuPAGE Bis-Tris gel (#NP0336BOX, Thermo Fisher) at 175V for 12 min, and stained as above. Gels were diced and washed repeatedly in 25 mM triethylammonium bicarbonate (TEAB, #T7408, Sigma-Aldrich) in 50 v/v % acetonitrile (ACN, #34998, Sigma-Aldrich) followed by 100% ACN. Gel pieces were then incubated in 25 mM TEAB and 10 mM dithiothreitol (#D11000, Research Products International) at 57°C for 1 hour, followed by iodoacetamide (#I6125, Sigma-Aldrich) at room temperature for 45 min. Gel pieces were dehydrated stepwise with 25 mM TEAB, 25 mM TEAB in 50% ACN and 100% ACN. After overnight digestion at 37°C with trypsin (#V5111, Promega), reaction was quenched with formic acid, and supernatant was collected. The remaining gel pieces were washed with 25 mM TEAB followed by 100% ACN, with both supernatants collected and pooled together. Pooled supernatants were dried in speed vacuum with heating and dissolved in 100 mM TEAB supplemented with formaldehyde and sodium cyanoborohydride. For dimethyl labelling, different samples were either labeled with light (CH_2_O, #252549, Sigma-Aldrich) or medium (CD_2_O, #sc-228228, Santa Cruz Biotechnology) formaldehyde. Corresponding light- and medium-labeled samples were subjected to electrospray ionization mass spectrometry on an Orbitrap Fusion Lumos Mass Spectrometer (Thermo Fisher). Results were obtained by searching against the *E. coli* strain K12 peptide database (Supplementary dataset 1).

### GO term comparison with P-bodies/stress granules

We identified human proteins that are components of P-bodies and/or stress granules based on “Gold standard” entries in the RNA granule database (https://rnagranuledb.lunenfeld.ca/). GO terms were assigned to the (*E. coli*) HP body, (human) P-body, and (human) stress granule proteins based on annotations from UniProt. We then used custom-written python code to identify the sets of GO terms present at the two-way and three-way interfaces of the GO term lists. We assessed significance via a permutation test in which we simulated the null distribution by randomly reassigning the set of *E. coli* K12 proteins assigned to HP bodies, and calculating the GO term overlaps with P-bodies and/or stress granules for each of 1000 permutations.

### Mammalian cell culture, plasmids and transfection

HEK293 cells (ATCC® CRL-1573™, ATTC, Manassas, VA, USA), HeLa cells (ATCC® CCL-2™, ATTC, Manassas, VA, USA) and NIH3T3 cells (ATCC® CRL-1648™, ATTC, Manassas, VA, USA) were grown and maintained in DMEM (Gibco™ 11995065) supplemented with 10% w/v Fetal Bovine Serum (#F4135, Sigma-Aldrich) and 1% w/v Penicillin-Streptomycin (#SV30010, Cytiva). All cells were cultured in a 37°C incubator at 5% CO_2_. EcPPK1 was a gift from Michael Downey (Addgene plasmid # 108850). Transfections were carried out on cells seeded on glass coverslips in a 24-well plate using Lipofectamine™ LTX Reagent with PLUS™ Reagent (#15338100, ThermoFisher Scientific) as per the protocol provided on the manufacturer’s website. Cells were transfected for 24 hours prior to treatment with sodium arsenite.

### Generation of yPPX expressing stable cell lines

yPPX fused to a 3xFLAG tag on the N-terminus was cloned into the pLVX-pTuner Green vector (# 632176, Takara Bio). The vector contained the coding sequence for an additional N-terminal destabilization domain (DD), which causes the degradation of the protein in the absence of the small molecule Shield1 (#632189, Takara Bio). Lentiviruses were generated with this construct at The Vector Core (University of Michigan Medical School). NIH3T3 cells were transfected with the DD-3x-FLAG-yPPX lentivirus in the presence of polybrene (#TR-1003-G, EMD Millipore) at a concentration of 10µg/ml. Cells were transfected for 24 hours and allowed to recover for 24 hours after removal of transfection media. Positively transfected clones were sorted based on ZsGreen fluorescence with the Cytoflex SRT (Beckman Coulter) using the B525 channel. Sorted cells were grown in the presence or absence of 0.5µM Shield1. The expression of yPPX was verified by western Blotting using a mouse anti-FLAG M2 antibody (#F3165, Sigma-Aldrich).

### Mammalian cell immunofluorescence staining and microscopy

To visualize P-bodies and stress granules via fluorescence microscopy, transfected cells were detached using 0.25% w/v Trypsin-EDTA (#T4049, Sigma-Aldrich) and seeded onto 12 mm circular coverslips (#72230-01, Electron Microscopy Sciences) placed in a 24-well plate at a density of 50,000 cells/well. The cells were treated with buffer or 0.5 mM sodium arsenite (#S7400, Sigma-Aldrich) for 30 mins. Afterwards, the cells were fixed with freshly prepared 4% v/v paraformaldehyde (#1578100, Electron Microscopy Sciences) for 20 mins. Fixed cells were washed 3 times with Phosphate Buffered Saline (PBS) before permeabilization with 0.3% v/v Triton X-100 (#T8787, Sigma-Aldrich) for 1 hour. Triton X-100 was prepared in a solution of 1% w/v BSA (#A3059, Sigma-Aldrich) in PBS. After permeabilization, cells were washed with PBS and incubated in blocking solution (1% w/v BSA in PBS) for 1 hour. To visualize endogenous polyP, cells were incubated with PPXBD-mCherry at a concentration of 10 μg/ml prepared in blocking solution overnight at room temperature. To stain P-bodies, a mouse monoclonal antibody against EDC4 (#sc-374211, Santa Cruz Biotechnology) or a rabbit monoclonal antibody against DCP1A (# NBP2-59785, Novus Biologicals) were used at a concentration of 2 μg/ml. To stain stress granules, a rabbit polyclonal antibody against G3BP1 (#130572AP, Proteintech) was used at 1 μg/ml. Cells were incubated simultaneously with the PPXBD-mCherry probe and the respective primary antibodies. The next day, cells were washed with 1X PBS and incubated with respective secondary antibodies for 2 hours at room temperature protected from light. The secondary antibodies that we used were goat anti-mouse IgG-Alexa Fluor® 647 (#A21235, Invitrogen) and goat anti-rabbit IgG-Alexa Fluor® 488 (#A11008, Invitrogen). Cells were then washed with 1X PBS and incubated with DAPI (#D1306, Thermo Fisher Scientific) at a concentration of 1 μg/ml for 10 mins to stain the nucleus. Cells were washed 3 times before mounting them onto a microscope objective slides using Prolong Gold (#9071S, Cell Signaling Technology) as the mounting medium. Mounted coverslips were sealed with nail polish and imaged 24 hours post mounting. Mounted cells were visualized using a 63x oil objective on a Leica SP8 laser scanning confocal microscope (Leica GmbH, Mannheim Germany) on a DMI8 base using the LAS X software.

